# Standardised method for cardiomyocyte isolation and purification from individual murine neonatal, infant, and adult hearts

**DOI:** 10.1101/2021.09.30.462562

**Authors:** Amy M Nicks, Sara R Holman, Andrea Y Chan, Michael Tsang, Paul E Young, David T Humphreys, Nawazish Naqvi, Ahsan Husain, Ming Li, Nicola J Smith, Siiri E Iismaa, Robert M Graham

**Affiliations:** Victor Chang Cardiac Research Institute, Darlinghurst, NSW 2010, Australia; St Vincent’s Clinical School, University of NSW, Kensington, NSW 2052, Australia; Department of Medicine (Cardiology), Emory University School of Medicine, Atlanta, GA, USA

**Author notes:** Corresponding authors: Robert M Graham or Siiri E Iismaa, Victor Chang Cardiac Research Institute, 405 Liverpool Street, Darlinghurst, NSW 2010, Australia, or. Cardiac Regeneration Research Institute, Wenzhou Medical University, Wenzhou, 325035, China (Ming Li).

**Keywords:** cardiomyocyte biology, cardiomyocyte isolation, postnatal heart development, cardiac hypertrophy, cardiomyocyte maturation, sex differences

## Abstract

Primary cardiomyocytes are invaluable for understanding postnatal heart development. However, a universal method to obtain freshly purified cardiomyocytes without using different age-dependent isolation procedures and cell culture, is lacking. Here, we report the development of a standardised method that allows rapid isolation and purification of high-quality cardiomyocytes from individual neonatal through to adult C57BL/6J murine hearts. Langendorff retrograde perfusion, which is currently limited to adult hearts, was adapted for use in neonatal and infant hearts by developing an easier *in situ* aortic cannulation technique. Tissue digestion conditions were optimised to achieve efficient digestion of hearts of all ages in a comparable timeframe (<14 min). This resulted in a high yield (1.56-2.2×10^6^ cells/heart) and viability (~70-100%) of cardiomyocytes post-isolation. An immunomagnetic cell separation step was then applied to yield highly purified cardiomyocytes (~95%) as confirmed by immunocytochemistry, flow cytometry, and qRT-PCR. For cell-type specific studies, cardiomyocyte DNA, RNA, and protein could be extracted in sufficient yields to conduct molecular experiments. We generated transcriptomic datasets for neonatal cardiomyocytes from individual hearts, for the first time, which revealed nine sex-specific genes (FDR<0.05) encoded on the sex chromosomes. Finally, we also developed an *in situ* fixation protocol that preserved the native cytoarchitecture of cardiomyocytes (~94% rodshaped post-isolation), and used it to evaluate cell morphology during cardiomyocyte maturation, as well as capture spindle-shaped neonatal cells undergoing cytokinesis. Together, these procedures allow molecular and morphological profiling of high-quality cardiomyocytes from individual hearts of any postnatal age.

## 1 Introduction

In mammals, postnatal heart development is characterised by a remarkable increase in mass that is required for normal cardiovascular responses in the adult animal. There are a variety of intrauterine growth factors that influence this cardiac growth, including hypoxia, nutrition, premature birth, endocrine stress, and congenital heart diseases[1–3]. However, the molecular events involved in normal cardiomyocyte maturation after birth are not well understood, including the pathological consequences of acquired heart diseases. The postnatal period also represents a therapeutic opportunity for the treatment of many types of cardiovascular diseases in children and adults[4–8].

Model systems for studying postnatal cardiomyocyte development are lacking, particularly due to the very limited availability of human samples that span different ages, whilst *in vitro* cell lines are not presently adequate for this purpose[9]. Investigations of cardiomyocyte ontogeny and disease-related alterations therein, is best achieved using primary murine cardiomyocytes[9], if they can be readily isolated at any developmental age. To date, there is no established universal method for the isolation and purification of cardiomyocytes from neonatal as well as adult hearts or ages in-between. Isolating neonatal or adult rodent cardiomyocytes from the cardiac milieu is not trivial and methods have been continually modified and improved since the 1960’s[10,11]. Langendorff retrograde perfusion is the gold standard for isolating high quality *adult* cardiomyocytes[12], but has also been used to isolate cardiomyocytes from 21 day-old mouse hearts[13]. To generate isolated *neonatal* cardiomyocytes, more rudimentary “batch chopping” procedures are used that involve crude mincing of hearts followed by sequential enzymatic digestion steps or an overnight digestion[12,14]. This protocol is suboptimal as it requires pooling 5-15 hearts at postnatal day (P) 0-3 for a single replicate[12,14], thereby using a larger number of animals, and precludes the evaluation of individual hearts. Hence, current methods used to isolate cardiomyocytes are age-dependent and differ considerably with respect to processing times, tissue digestion conditions and number of hearts used per isolation.

Once adult cardiac cells are isolated after Langendorff perfusion and tissue disaggregation, cardiomyocytes are generally enriched by three rounds of differential centrifugation for 3 min[15] or by gravity settling for 15 min[16] to remove a proportion of non-myocytes, i.e., endothelial cells and fibroblasts. Cardiomyocytes may then be further purified by cell culture (<24 hr or up to 72 hr) or occasionally by Percoll gradient centrifugation[12,15,16]. Neonatal murine cardiomyocytes are generally purified, without enrichment, by differential cell culture that involves cardiac cells being pre-plated onto an uncoated culture dish for several hours to allow non-myocytes to attach. The supernatant, containing cardiomyocytes, is then transferred to a collagen-coated culture dish for 12 hours to 3 days[14,17]. This is problematic given the time-consuming incubation steps and nature of cell culture purification that leads to morphological and functional changes in cells, such as altered molecular composition (RNA, protein, metabolites), abnormal cytoarchitecture, cardiomyocyte dedifferentiation, and cell death[12,14].

Here, we developed a standardised Langendorff perfusion and immunomagnetic cell separation procedure to rapidly isolate and purify cardiomyocytes in high yield and viability using individual neonatal, infant, and adult murine hearts. We describe, for the first time, sex differences in the neonatal cardiomyocyte transcriptome. A further advance was the use of an *in situ* fixation protocol to assess cardiomyocyte morphology in terms of cell growth and binucleation during postnatal development, and to capture isolated neonatal cells in mitosis. Application of these techniques should provide further important insights into cardiomyocyte biology and heart development.

## 2 Methods

A detailed description of the methods is available in the Supplementary material online.

### 2.1 Animals

C57BL/6J mice were used at various ages: both male and female neonates (postnatal day 0-2, P0-2) and infants (P10 and P13); and male P70 adults (9-11-week-old). Animals were housed in individually ventilated cages, given food and water *ad libitum*, and exposed to a 12-hr light-dark cycle. All studies were approved by the local Garvan Institute /St Vincent’s Hospital Animal Ethics Committee (No. 13/07, 16/16, 19/17) and performed in accordance with The Australian Code of Practice for the Care and Use of Animals for Scientific Purposes (2021).

### 2.2 *In situ* aortic cannulation

Mice were administered an intraperitoneal injection of heparin for 20-30 min before being sacrificed using age-appropriate methods, as detailed in Table 1. After sterilizing the chest with 70% ethanol, the anterior chest wall (ribcage) was removed for *in situ* cannulation of the aorta (described in Fig. 1) in preparation for Langendorff perfusion of the heart. This was performed using age-specific microsurgical instruments and materials (Fig. 2A, Fig. 2B and Supplementary Table S1). This method of cannulation uses a longer length of aorta (~0.5-1 cm) that is easier to identify and guide onto the cannula. Given the marked differences in aortic diameter and heart size at different postnatal ages (Fig. 2C), needles of appropriate sizes were required for the aortic cannulations (Fig. 2D and Table 1). With these adaptations, even the smaller diameter and more fragile neonatal and infant aortae could be cannulated rapidly (~3 min) under a dissection microscope. Adult hearts were cannulated within a minute and a microscope was only used to secure and confirm the correct positioning of the cannula.

**Fig. 1.**
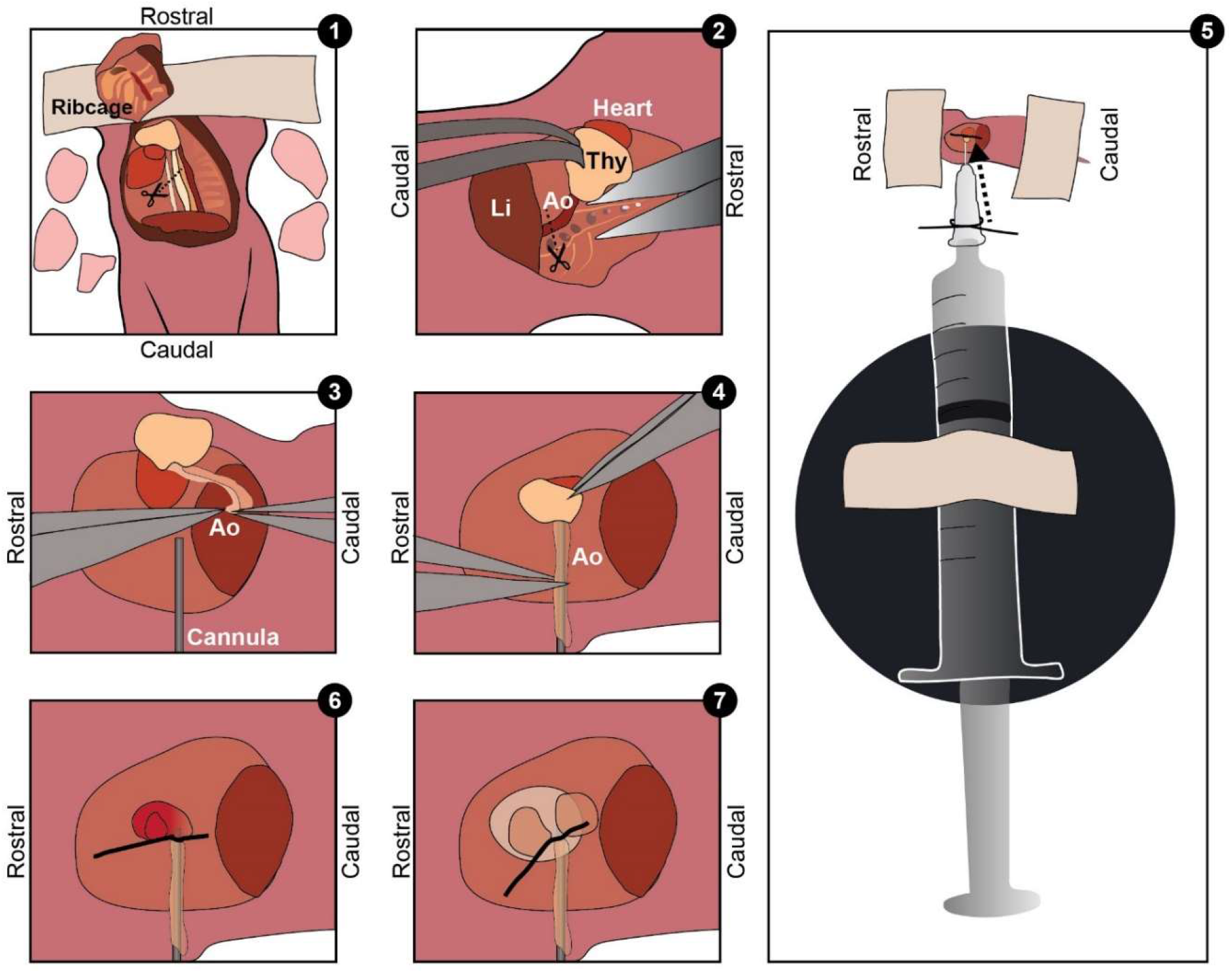
*In situ* cannulation of the neonatal murine heart. *Step 1:* The ribcage of a neonatal C57BL/6J mouse (postnatal day 2, P2) is opened and the five lobes of the lungs removed, followed by the transection of the inferior vena cava and oesophagus (see scissor line). *Step 2:* The operating board is rotated 90° (right-handed operator; liver [Li] is left). To release the aorta from the spinal cord, fine-forceps are used to lift the heart via the attached thymus (Thy) and fine-curved scissors (in the dominant hand) are used to cut the connective tissues surrounding the heart, releasing it. Then the thymus and heart are elevated whilst the scissors are pressed flat against the spine to carefully dissect the connective tissue attaching the aorta (Ao) to the spine, starting from the aortic arch down to the diaphragm, where the aorta is transected (see scissor line). Note: This step results in the isolation of the thymus-heart-aorta *en bloc;* the heart then being ready for cannulation. *Step 3 and 4:* The operating board is rotated 180°. With the heart resting *in situ*, the aorta, now drained of blood, is drawn onto the cannula (a blunted 30G needle attached to a syringe) by gripping the edges of the vessel walls with fine-forceps and finding the opening, which is pulled onto the cannula until the tip of the cannula is just above the aortic valve. *Step 5:* The set-up of the cannula attached to the syringe (10 mL), pre-filled with perfusion buffer, is shown relative to the size of a P2 mouse. A 6-0 silk suture tied loosely around the base of the cannula, prior to cannulation, is re-positioned around the cannulated aorta between the thymus and atria, and then secured using a double knot. Step *6:* The thymus is removed prior to mounting the heart onto a Langendorff perfusion rig for perfusion. Step *7:* during practice cannulations, the heart can be flushed with perfusion buffer using the attached syringe. Blanching and expansion of the heart confirms the correct positioning of the cannula. Note: neonates and infants have smaller, shorter, and more delicate aortae that must be handled gently to avoid to breakage.

**Fig. 2.**
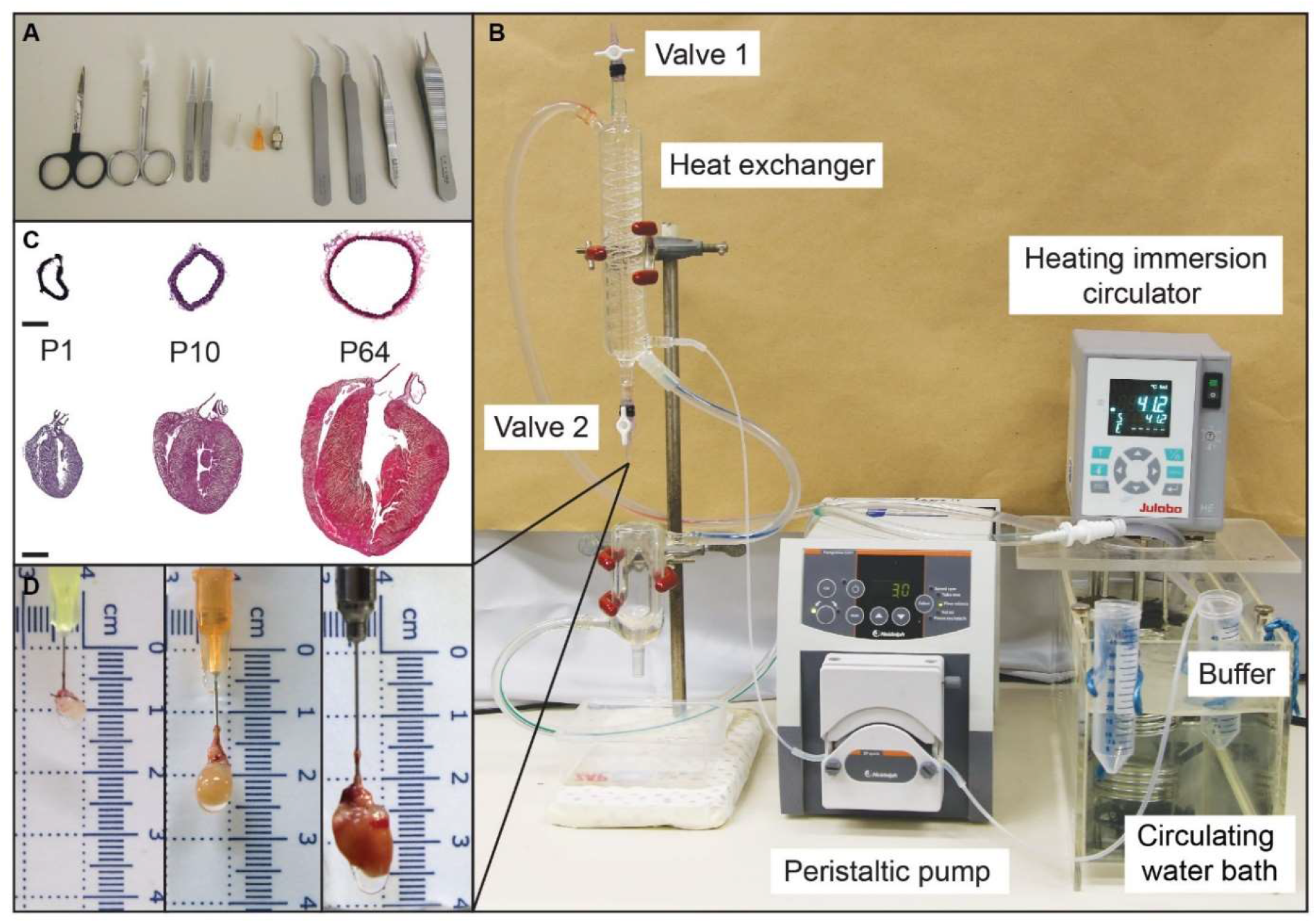
Instruments and apparatus used for *in situ* aortic cannulation and Langendorff perfusion of murine hearts. (A) Microsurgical tools from left to right: tough-cut straight sharp scissors, fine-sharp scissors, mini forceps x2, blunted 30G and 25G needles, and 24G gavage needle, curved forceps x2, and curved and straight serrated forceps. (B) Langendorff apparatus consisting of a heated immersion circulator to control water bath temperature. The heated water is circulated through the insulating jacket of the heat exchanger via a peristaltic pump (John Morris Group, Australia) that sends buffer through the tubing and internal coil of the heat exchanger allowing tissue perfusion at a regulated flow rate and temperature. Immediately before mounting the heart, a pressure gradient is created by closing valve 1 (in the horizontal position), opening valve 2 (in the vertical position), and turning on the pump. The water-jacketed insulating organ bath is moved upwards to surround the heart and maintain a warm environment. (C) H & E stained (DNA, blue; and cell cytoplasm, red) aortae sectioned at the level of the diaphragm (scale bar is 200 μm), and coronal heart sections (scale bar is 1 mm) from C57BL/6J neonatal (P1), infant (P10) and adult (P64) mice. (D) P1, P10 and P70 (left to right) cannulated murine hearts mounted onto the Langendorff apparatus. Size differences shown by the background scales.

**Table 1:** Cardiomyocyte (CM) isolation and purifcation details.

|  | Neonate (P0-P3) | Infant (P10-13) | Adult (P70) |
| --- | --- | --- | --- |
| **Pre-cannulation** |  |  |  |
| Heparin (1000IU/mL) | 10 µl | 20 µl | 100 µl |
| Incubation time post-heparin (min) | 20-30 | 20-30 | 20-30 |
| **Cannulation step** |  |  |  |
| Euthanasia | Decapitation | Decapitation (≤P10) or cervical dislocation (>P10) | 2-4% Isoflurane and cervical dislocation |
| Cannula | 30G blunt needle | 25G blunt needle | 24G gavage needle |
| Heart and aorta dissection, and cannulation time (min) | 5 | 5 | 5 |
| **Perfusion step** |  |  |  |
| Flow rate (mL/min) | 1.5 | 2 | 3 |
| Perfusion buffer volume (mL) | 4 | 6 | 9 |
| Perfusion time with perfusion buffer (min:sec) | 2:40 | 3 | 3 |
| Weight for enzyme conc. (g) | 5 | 8 | Actual body weight |
| Digestion buffer volume (mL) | 16 | 16-20 | 25-30 |
| Perfusion time with digestion buffer (min:sec) | 10:40 | 8-10 | 8:20-10 |
| **CM isolation step** |  |  |  |
| Buffer | Transfer buffer | Transfer buffer | Transfer buffer |
| Pipette | P1000 | 5.8 mL transfer pipette | 7.7 mL transfer pipette |
| Filter (µm) | 70 | 200 | 200 |
| Cell isolation time (min) | 10 | 10 | 10 |
| **CM enrichment step** |  |  |  |
| Centrifugation | NA | 55 g for 3 min (x3) | 20 g for 3 min (x3) |
| Buffer volume (mL) | NA | 8 | 10 |
| **CM purification step** |  |  |  |
| Method | Miltenyi Neonatal CM Isolation kit | CD31 Ab conjugated to Dynabeads | CD31 Ab conjugated to Dynabeads |
| Ab-bead volume (µl) | 10 | 12.5 | 25 |
| Ab-bead incubation time (min) | 15 | 15 | 15 |
| Total time (min) | 50 | 60 | 60 |

P, postnatal day; Ab, antibody.

### 2.3 Langendorff retrograde perfusion

Using the Langendorff apparatus (Fig. 2B), perfusate was maintained at 37°C by adjusting the water bath temperature and checking the outflow temperature at valve 2 using a thermometer probe (cat. #: BAT-10, Physitemp). Once the heart was mounted onto the apparatus, it was flushed of blood using a modified Krebs-Ringer solution (without Ca^2+^) referred to as “perfusion” buffer and then perfused with “digestion” buffer that incorporates proteolytic enzymes (collagenase B, D, and protease XIV; buffer composition and recipe in Supplementary Table S2, S3). The conditions for efficient tissue digestion were optimised for neonatal through to adult hearts (<14 min), including flow rate, concentrations of proteolytic enzymes, and volumes of buffers (details in Table 1). After digestion, the atria were removed and the cardiac ventricles were collected in a petri dish containing transfer buffer (composition in Supplementary Table S2). The methods describing Langendorff perfusion and tissue digestion can be found in the Supplementary methods online.

### 2.4 Cardiomyocyte isolation and enrichment

Cardiomyocytes were isolated from the ventricles (neonatal, infant and adult) by disaggregation and gentle trituration of the digested heart tissues, which was then passed through an appropriately sized cell filter cup (Table 1) to remove undigested tissues and collect cells. Adult (P70) cardiomyocytes, which are much larger than non-myocytes, were enriched by differential centrifugation (Table 1). This step was adapted for infant (P10 and P13) cardiomyocytes, which are smaller than adult cells, by increasing the centrifugal force to minimise cardiomyocyte losses. Detailed protocols are given in the Supplementary methods online.

### 2.5 Cardiomyocyte purification

Infant and adult cardiomyocytes were purified by immunomagnetic bead-based cell depletion of endothelial cells that were the main contaminating non-myocyte population (materials listed in Supplementary Table S1). Rat anti-mouse CD31 antibody was mixed with sheep anti-rat IgG Dynabeads in “isolation” buffer and incubated overnight with rotation at 4°C (preparation details in Supplementary Table S2, S4). The resulting Ab-conjugated beads (Ab-beads) were stored at 4°C. Ab-beads (25 or 50 μl per infant and adult sample, respectively) were prepared before use by resuspending and washing three times (1 mL transfer buffer) using a MagnaRack (Thermo Fisher) to remove excess unbound antibody (details in Table 1). To purify enriched fractions of cardiomyocytes, samples were resuspended in transfer buffer (2 mL) and added to the washed Ab-beads and mixed on a roller (15 min, RT). Each sample was then placed onto the MagnaRack (2 min) to separate bead-bound endothelial cells from unbound cardiomyocytes. To ensure complete removal of beads, the supernatant fraction was transferred, using uncut transfer pipettes, consecutively into two fresh 2 mL tubes on the MagnaRack and the supernatant was finally collected into a 15 mL Falcon tube. Two additional gentle bead washes of the original tube (2 mL transfer buffer per wash) were similarly processed to increase cardiomyocyte yield and were also collected into the 15 mL Falcon tube (total of 6 mL). Cardiomyocytes were gently resuspended before cell counting using a transfer pipette (7.7 mL) in either 6 ml or 8ml of transfer buffer for infant or adult cardiomyocyte preparations, respectively. Cardiomyocytes were counted using a haemocytometer and the proportion of viable, rod-shaped cells determined.

Neonatal cardiomyocytes were purified by immunomagnetic separation using the Neonatal Cardiomyocyte Isolation Kit (Miltenyi Biotec), according to manufacturer’s instructions. The protocol was performed using transfer buffer instead of the kit prescribed isolation buffer. Briefly, cardiac cell suspensions were centrifuged (300 *g*, 5 min), and the supernatant removed prior to incubation (15 min, 4°C) with magnetic MicroBeads bound to various antibodies (manufacturer’s proprietary information) directed at non-myocyte cell surface antigens. The mixture was resuspended in transfer buffer (1.5 mL) and passed through a magnetically-charged iron matrix column having been placed on a magnetic rack (Supplementary Table S1) and the eluate containing cardiomyocytes was collected. The column was washed twice with transfer buffer (1.5 mL) and eluates collected and pooled. After centrifugation (300 *g*, 5 min), cardiomyocytes were gently resuspended in transfer buffer (3 mL) using a transfer pipette (5.8 mL) and counted on a haemocytometer. Cell viability was almost 100% with round neonatal cardiomyocytes also excluding Trypan blue.

Overall, the time taken from aortic cannulation to complete cardiomyocyte purification was less than one hour for all postnatal ages. Cell pellets were stored at −80°C, or fixed post -isolation, -enrichment or -purification in 2% paraformaldehyde (PFA, 5 min, RT) followed by wash steps (2 × 10 mL PBS, 100 g-600 g, 3 min, RT) and then stored in PBS at 4°C for subsequent experiments.

### 2.6 Analyses of cardiomyocyte purity

Cardiomyocyte purity was assessed using three different methods: immunocytochemistry, flow cytometry and qRT-PCR (detailed protocols are available in the Supplementary methods online). Briefly, for immunocytochemistry experiments using confocal microscopy, cells were stained for cardiomyocytes (cardiac troponin T, cTnT), endothelial cells (isolectin B4, IB4), and nuclei (TO-PRO-3). Flow cytometric evaluations were performed by the intracellular staining of cardiomyocytes using cTnT and nucleated cells were identified by propidium iodide (PI). Standard techniques were used for cDNA synthesis and qRT-PCR to measure the expression of cell marker genes (TaqMan probes listed in Supplementary Table S5).

### 2.7 Cardiomyocyte DNA, RNA, and protein yields

DNA, RNA and protein extraction methods are detailed in the Supplementary methods online.

### 2.8 RNA-Sequencing of neonatal cardiomyocytes

Purified cardiomyocytes from individual male and female neonatal (P2) hearts were used to extract poly(A) RNA for sequencing using a service provider (Australian Genome Research Facility, Melbourne, Australia). RNA-Seq analyses were performed as detailed in the Supplementary material online. Transcriptomic datasets were deposited on the Gene Expression Omnibus database with the accession number: GSE193137.

### 2.9 In *situ* fixation of cardiac cells

Cardiac cells were fixed *in situ* immediately after Langendorff perfusion by flushing the digested heart with 2% PFA (2-5 mL) using a 5 mL syringe. Atria were removed and the heart was submerged in 2% PFA (5 min). Tissue was disaggregated and passed through an appropriately-sized cell filter cup (details in Table 1 and Supplementary material online). Cell suspensions were further fixed (2% PFA, 5 min, RT), washed (2x 10 mL PBS, 100-600 *g*, 3 min) and stored in PBS at 4°C. The total time taken for *in situ* fixation was 30 min per heart.

### 2.10 Statistical analysis

Data are presented as the mean ±SD unless otherwise stated. Statistical tests were performed using one-way ANOVA combined with Dunnett’s multiple comparisons, or unpaired Welch’s t-test if variance was unequal between two groups, or, correlation tests performed using Pearson’s correlation coefficient. GraphPad Prism 9 (v9.1.2) was used for statistical analyses and presenting graphical data. Differences were considered statistically significant if P<0.05.

## 3 Results

### 3.1 Isolation of high-quality cardiomyocytes from neonatal and infant hearts

The cardiomyocyte and purification procedure developed here consists of five steps resulting in purified cells within one hour at all postnatal ages (Fig. 3). The first step was the cannulation of the heart in preparation for Langendorff perfusion. Typically, adult hearts are placed into a petri dish of ice-cold buffer and cannulated *ex vivo* using a short section of the ascending aorta (transected before or at the aortic arch)[15,16], however, this can be technically challenging and time-consuming particularly for infant and neonatal hearts. To obviate this, we performed all cannulations *in situ*, i.e., with the heart remaining in the thoracic cavity, and used a longer length of the descending aorta (transected at the diaphragm), which made it easier to identify and guide the aorta onto the cannula (Fig. 2).

**Fig. 3.**
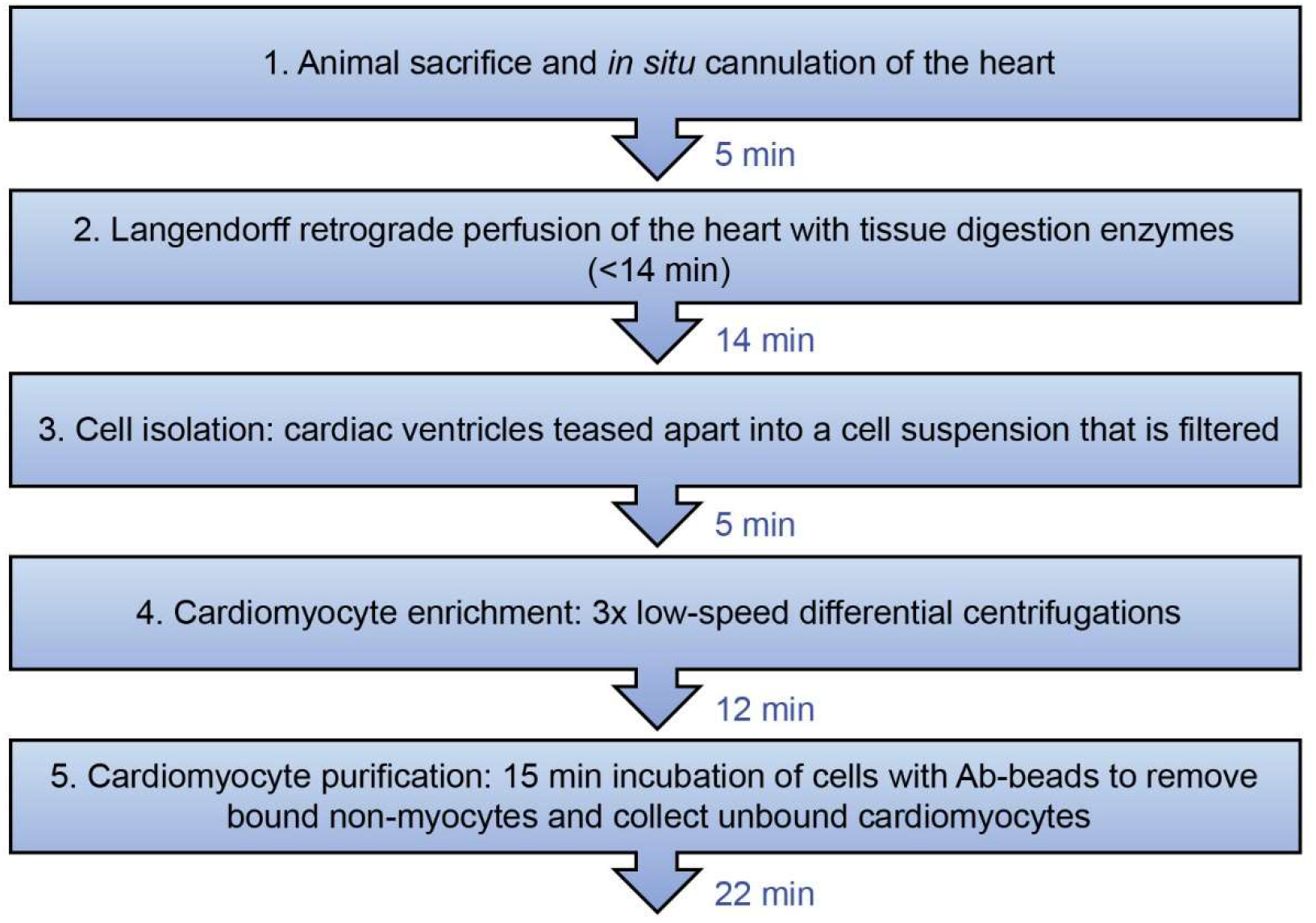
Overview of the cardiomyocyte isolation and purification procedure (<1 hr). The five steps required for cardiomyocyte purification are shown with the estimated time taken per step. For *in situ* fixation of cardiac cells, the heart was perfused with PFA immediately after step 2. Cardiomyocyte enrichment (step 4) was omitted for neonatal cardiac cells. See Methods for further details.

In the second step (Fig. 3), Langendorff perfusion was adapted and optimised for the digestion of neonatal, infant, and adult hearts by choosing effective proteolytic enzymes, adjusting enzyme concentrations and buffer volumes, as well as perfusion flow rates (Table 1). These conditions were optimised to standardise the time taken for tissue digestion across all ages (<14 min). After the third step of cardiomyocyte isolation by tissue disaggregation and cell filtration (Fig. 3), cardiomyocytes were isolated in high yield from P2 hearts (~1.56×10^6^ cells/heart), P10 or P13 hearts (~1.62×10^6^ cells/heart), and P70 hearts (~2.21×10^6^ cells/heart) (Table 2). The viability of cells, assessed based on the number of rod-shaped vs round infant and adult cardiomyocytes, was ~72% (ranging from 60-90%, Table 2). “Batch chopping” methods yield rounded neonatal cardiomyocytes[14]. It was surprising, therefore, to find that more than half of the neonatal cardiomyocytes we isolated were rod-shaped (~53%, ranging from 30-90%, Table 2). The shape of neonatal cardiomyocytes is not indicative of cell viability, almost all cells, including rounded cells, were viable and excluded Trypan blue dye.

**Table 2:** Cardiomyocyte (CM) yield after the isolation and purification steps.

| Age | CM number after isolation | Viability (%) | Rod-shaped (%) | n | CM number after purification | n |
| --- | --- | --- | --- | --- | --- | --- |
| Neonate (P2) | 1.56±0.48x10<sup>6</sup> | 100 | 53 (~30-85) | 72 | 1.03±0.33x10<sup>6</sup> | 56 |
| Infant (P10) | 1.62±0.39x10<sup>6</sup> | 70 (~60-90) | 70 (~60-90) | 69 | 0.96±0.29x10<sup>6</sup> | 44 |
| Infant (P13) | 1.65±0.44x10<sup>6</sup> | 70 (~60-90) | 70 (~60-90) | 25 | 0.8±0.21x10<sup>6</sup> | 25 |
| Adult (P70) | 2.21±0.36x10<sup>6</sup> | 72 (~60-90) | 72 (~60-90) | 27 | 1.6±0.29x10<sup>6</sup> | 19 |

Mean ±SD; P, postnatal day. Rod-shaped vs round-shaped CMs (%).

### 3.2 In *situ* fixed cardiomyocytes retained their *in vivo* cytoarchitecture

We developed a method for fixing cardiomyocytes *in situ* by PFA perfusion of the heart immediately after Langendorff perfusion with proteolytic enzymes. This led to ~94% of isolated cardiomyocytes being rod-shaped (or spindle-shaped in the case of neonatal cardiomyocytes). The proportion of spindle-shaped neonatal, and rod-shaped infant and adult cells after *in situ* fixation was ~43%, ~16% and ~30% higher, respectively, than that observed if cells were PFA fixed only post-isolation (Fig. 4A).

**Fig. 4.**
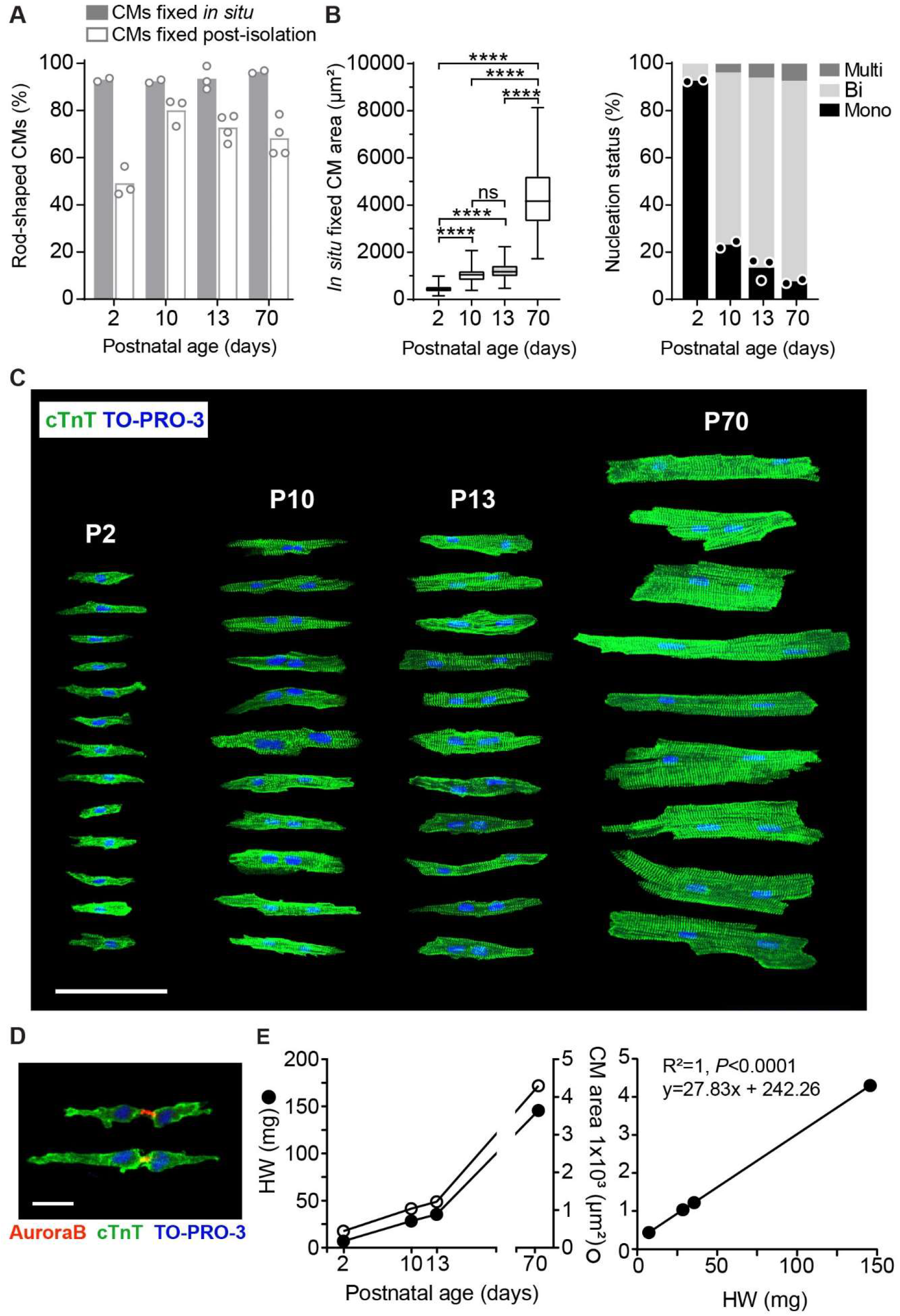
Gross morphology of maturing postnatal cardiomyocytes. (A) Percentage of rod-shaped vs round C57BL/6J ventricular cardiomyocytes with *in situ* fixation versus post-isolation fixation of neonatal (P2), infant (P10 and P13), and adult (P70) preparations (n=2-4, 120-420 cardiomyocytes). (B, left panel) Cardiomyocyte (CM) areas of *in situ* fixed cardiac cells determined from confocal immunofluorescence images of P2, P10, P13 and P70 cells (n=2-3, ~110-300 cardiomyocytes per age). Results are shown as box and whisker plots, and analysed by one-way ANOVA with Tukey’s multiple comparisons. ****P<0.0001. Right panel: percentages of mono-nucleated (1N), bi-nucleated (2N), or multi-nucleated (>2N) cardiomyocytes were counted (n=2-3, ~100-400 cardiomyocytes per age). (C) Representative images of *in situ* fixed cardiomyocytes at P2, P10, P13, and P70, immunostained with the cardiomyocyte-specific marker, cTnT (green), and nuclei with DNA dye, TO-PRO-3 (blue). Scale bar is 100 μm. (D) Neonatal (P2) *in situ* fixed cardiomyocytes showed Aurora B kinase staining between two daughter cells at the midbody (red). Scale bar is 20 μm. (E) Heart weight (HW, n=6-8, closed circles) and cardiomyocyte area (n=2-3, open circles) were compared throughout postnatal development at P2, P10, P13 and P70 (adult male) ages. Pearson’s correlation coefficient calculated for HW versus cardiomyocyte area.

### 3.3 Morphological assessment of postnatal cardiomyocyte maturation

Using cells fixed *in situ*, we characterised cardiomyocyte morphology during postnatal development. Cardiomyocytes at P2, P10, P13 and P70 were stained for cTnT (green) and nuclei (TO-PRO-3, blue) and cell area was measured by planimetry[18,19]. As expected, we observed considerable growth in cell size with cardiomyocyte maturation and found increases in cell area of 2.3-, 2.8- and 9.6-fold from P2 to P10, P13 and P70, respectively (Table 3, Fig. 4B). Moreover, changes in nucleation state during postnatal development were readily observed with *in* situ-fixed cells: 93% of P2 cardiomyocytes were mononucleated, whereas the majority (73-81%) became binucleated by P10-13 and thereafter; 85% being binucleated at P70 (Fig. 4B). *In situ* fixation showed that cardiomyocytes are small and spindle shaped at P2 and then enlarge and become more rod-shaped at P10, P13, and P70 (Fig. 4C). Cardiomyocytes are highly irregular in cell shape, having a jagged appearance of their intercalated discs at the cell edges (Fig. 4C). Not surprisingly, therefore, cell area was found to vary by up to a third at all ages (Table 3). Nevertheless, there was a very strong positive correlation (P<0.0001) between heart growth and cardiomyocyte area (Fig. 4E). *In situ* fixation also allowed isolated spindle-shaped neonatal cardiomyocytes to be captured undergoing cytokinesis, as evidenced by Aurora B kinase staining of the midbody and Flemming body (Fig. 4D).

**Table 3:** Cardiomyocyte characteristics, including DNA, RNA, and protein yield postpurification.

|  | Neonatal (P2) | Infant (P10) | Infant (P13) | Adult |
| --- | --- | --- | --- | --- |
| Binucleated (%) | 7 | 73 | 81 | 85 |
| Cell area ($\mu\text{m}^2$) | 446±148 | 1041±316 | 1226±333 | 4298±1278 |
| DNA yield (pg/cell) | 7.6 (2) | 13.8 (2) | nd | 12.1 (2) |
| RNA yield (pg/cell) | 7.3 (20) | 18.1 (12) | 23.5 (25) | 31.1 (12) |
| RIN score | 8.9 (20) | 9 (12) | 8.6 (25) | 8.5 (12) |
| Protein yield (pg/cell) | 105 (3) | 312 (3) | nd | 4026 (3) |

Mean ± SD; n number (); nd, not determined; RIN, RNA integrity number.

### 3.4 High cardiomyocyte yield after rapid purification by immunomagnetic cell separation

Infant and adult cardiomyocytes were enriched by three low-speed centrifugations to remove smaller non-myocytes (step 4, Fig. 3). This step was omitted for the enrichment of neonatal cardiomyocytes as their small size precludes efficient separation from non-myocytes. Despite the enrichment of infant and adult cardiomyocytes, endothelial cells were still present as undigested clusters or as strings that were sometimes attached to cardiomyocytes (Supplementary Fig. S1). We therefore implemented a purification step 5 (Fig. 3) using anti-CD31 antibody bound to Dynabeads (Ab-beads) to remove the contaminating endothelial cells. Neonatal cardiomyocytes were also purified after isolation by immunomagnetic bead-based cell separation using a kit (Neonatal Cardiomyocyte Isolation Kit, Miltenyi Biotec). Both purification steps were performed in a comparable timeframe across all ages (~22 min). After the enrichment step, total cardiomyocyte yield decreased by ~30% for P10 and P13 and by ~20% for P70 preparations, but cell viability remained unchanged. Despite some cell losses during enrichment and purification, the final yield of cardiomyocytes remained high (9×10^5^ per P2, P10, and P13 heart, and 1.5×10^6^ cardiomyocytes per P70 heart; Table 2). Importantly, the proportion of viable cardiomyocytes was unaltered after neonatal cardiomyocyte purification, while cell viability (rod-shaped) was either unaltered or only slightly reduced after infant and adult cardiomyocyte purification and structural integrity was maintained (Fig. 5A).

**Fig. 5.**
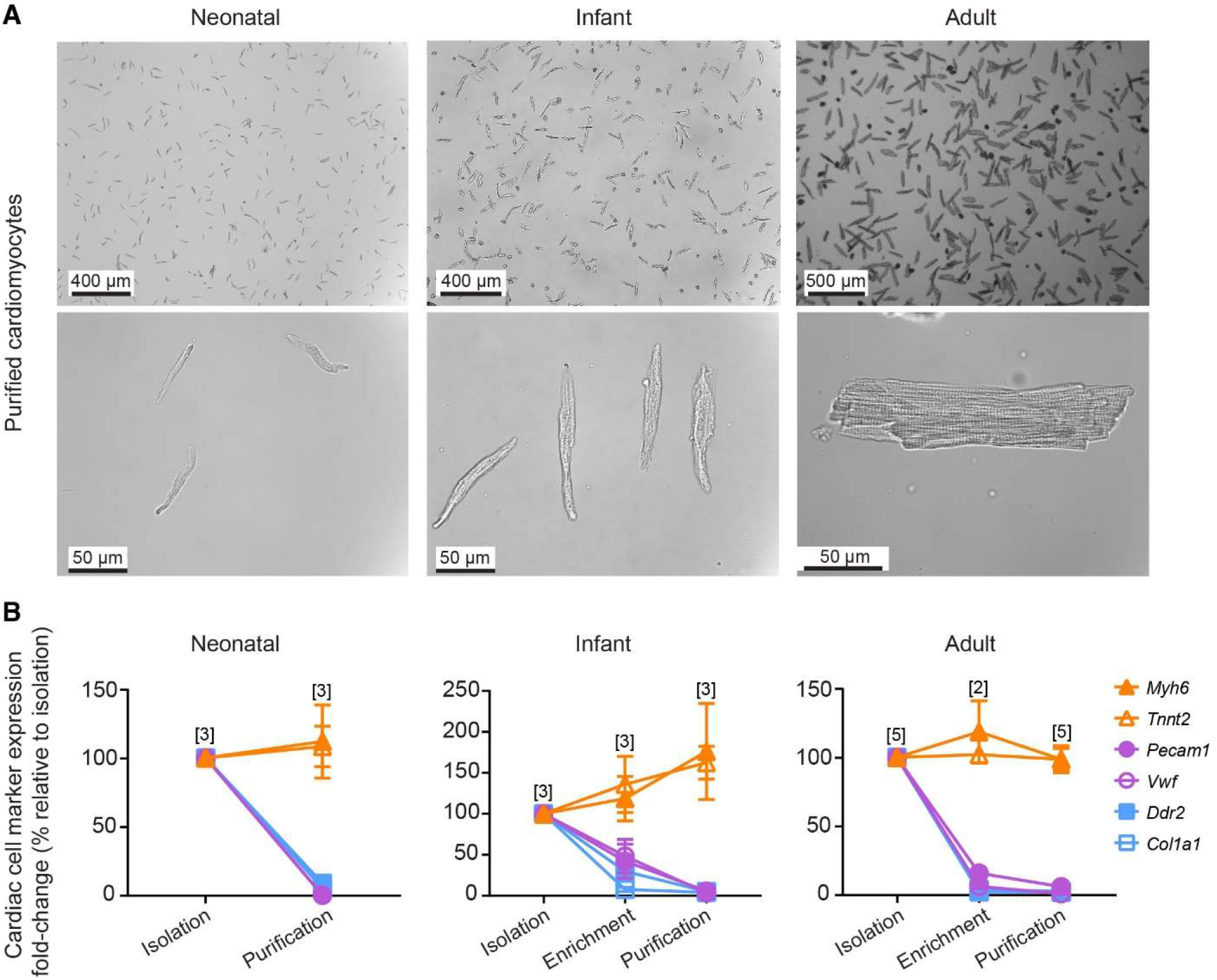
Cell morphology and cardiomyocyte purity post-purification. (A) Representative brightfield images of freshly purified high-quality cardiomyocytes (~80-85% rod-shaped CMs) from C57BL/6J neonatal (P2), infant (P10), and adult (8-10-week-old) mouse hearts. Cells were dispersed onto slides and images taken at 5X, 20X, 40X objectives on a Leica microscope and 4X on a Nikon Eclipse TS100 microscope. (B) Cardiomyocyte purity assessed by qRT-PCR gene expression levels of cardiac cell markers after cardiomyocyte isolation, enrichment and purification from: neonatal P2 (n=3); infant P10 (n=3); and adult P70 hearts (n=2-5). Genes were normalised to *Gapdh* and fold-change calculated relative to the isolation fraction using 2^−ΔΔCt^ method, shown as % (mean ±SEM). Cardiomyocyte markers (orange): *Myh6*, α-myosin heavy chain and *Tnnt2*, cardiac Troponin T; endothelial cell markers (purple): *Pecam1*, Platelet Endothelial Cell Adhesion Molecule 1, and *Vwf*, von Willebrand Factor; fibroblast markers (blue): *Ddr2*, Discoidin Domain Receptor 2 and *Col1a1*, Collagen Type I Alpha 1 Chain.

The cardiomyocyte DNA, RNA, and protein extracted from purified cells (Table 3) were in sufficient yield to conduct experiments using individual hearts. Moreover, the average RNA integrity number (RIN) score (~8.75 across all ages) indicated RNA of high-quality (Table 3). Altogether, the yield and quality of cardiomyocytes after the final purification step was high.

### 3.5 Expression of non-myocyte cell markers depleted after cardiomyocyte purification

To evaluate cardiomyocyte purity at the gene level, the expression of cardiac cell-type specific markers were assessed by qRT-PCR (Fig. 5B and Supplementary Table S6). This confirmed significant depletion of non-myocyte genes, including the endothelial cell markers, *Pecam1* (P2, 100% vs 0.37%, P<0.0001; P10, 100% vs 5.8%, P=0.0010; P70, 100% vs 6.17%, P<0.0001) and *Vwf* (P2, 100% vs 0.17%, P<0.0001; P10, 100% vs 3.39%, P=0.0003; P70, 100% vs 1.36%, P<0.0001), and fibroblast markers, *Ddr2* (P2, 100% vs 8.9%, P=0.0013; P10, 100% vs 5.51%, P<0.0001; P70, 100% vs 2.79%, P<0.0001) and *Col1a1*[20] (P2, 100% vs 4.98%, P=0.0002; P10, 100% vs 4.45%, P=0.0002; P70, 100% vs 2%, P<0.0001); all differences were determined using the unpaired Welch’s t-test. In contrast, the expression of cardiomyocyte cell markers, *Myh6* and *Tnnt2*, was not significantly different post-purification at any age.

### 3.6 Cardiomyocyte purity confirmed by immunocytochemistry and flow cytometry

Cardiac cell populations were quantified at each step of the cardiomyocyte purification procedure by immunocytochemistry using a cardiomyocyte-specific marker (cardiac troponin T, cTnT), endothelial cell marker (isolectin B4, IB4), and DNA intercalating dye to identify nuclei (TO-PRO-3; Fig. 6A and Supplementary Fig. S2-S5). “Other” cardiac cell types showed nuclear (TO-PRO-3^+^) but neither cTnT nor IB4 staining. This revealed that the differential centrifugation steps enriched the cardiomyocyte population (TO-PRO-3^+^, cTnT^+^) by +20%, +15% and +32% at P10, P13, and P70, respectively (Fig. 6B). The cardiomyocyte population significantly increased post-purification from 58% to 96% (P=0.0011) at P2, 36% to 96% (P<0.0001) at P10, 37% to 93% at P13 (P<0.0001), and 51% to 93% at P70 (P<0.0001, Fig. 6B), compared to the isolation fraction. The removal of endothelial cells was confirmed in the Ab-bead bound fraction that comprised 69%, 90%, 85% and 85% endothelial cells (TO-PRO-3^+^, IB4^+^) in P2, P10, P13 and P70 preparations, respectively (Fig. 6B). Cardiomyocyte purity was also assessed by flow cytometry (Fig. 6C), which confirmed significant increases in the cardiomyocyte population (PI^+^, cTnT^+^) from 82% to 99% in P2, 64% to 96% in P10, and 64% to 94% in P70 preparations after purification. Hence, immunomagnetic cell separation is an effective method to consistently purify cardiomyocytes.

**Fig. 6.**
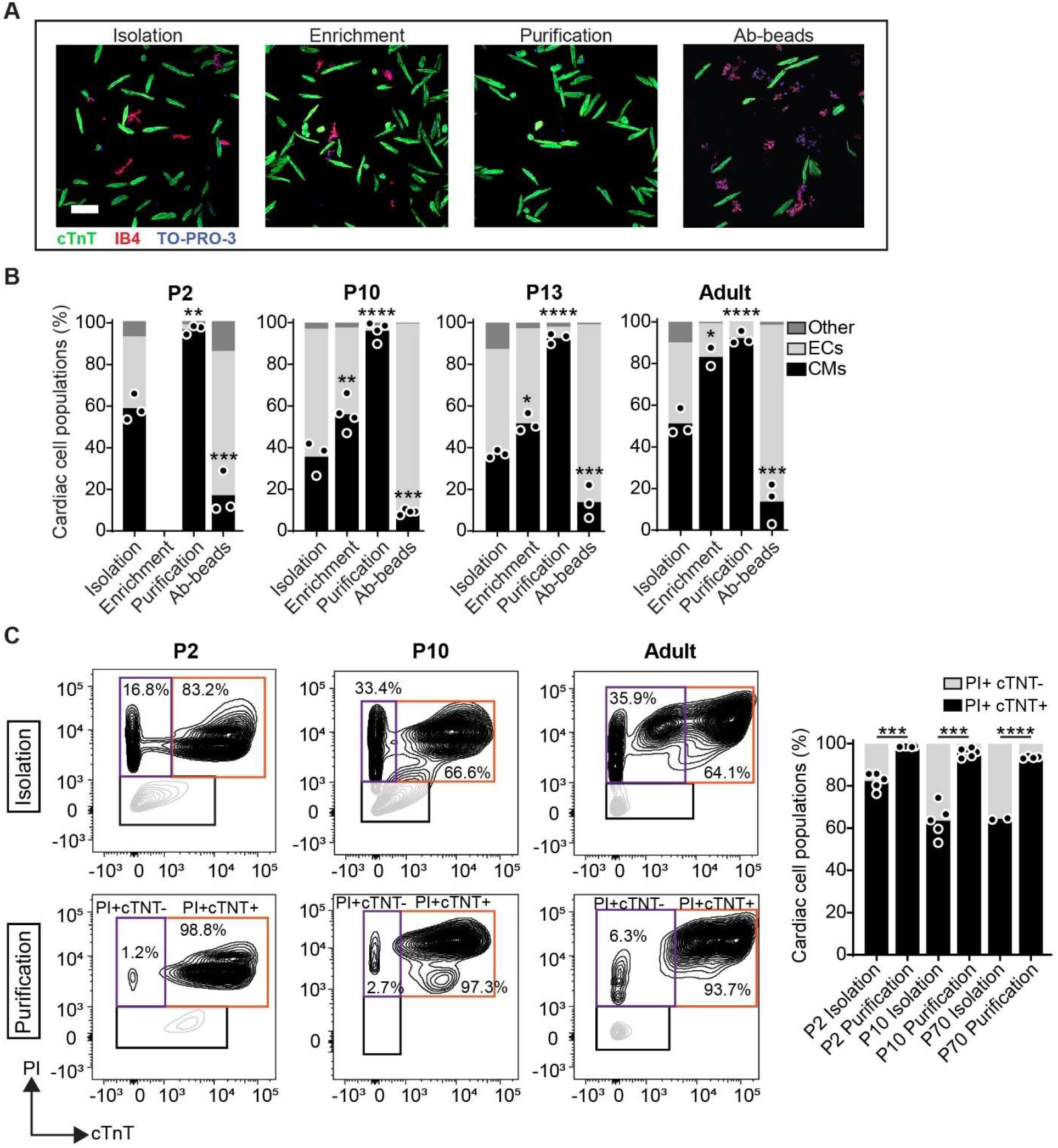
Cardiomyocyte purity assessed by immunocytochemistry and flow cytometry. (A) Representative immunofluorescence images of C57BL/6J P10 murine cardiac cells were stained to identify nucleated (DNA, TO-PRO-3, blue) cells, and co-stained with cardiac troponin T (cTnT, green) for cardiomyocytes or isolectin B4 (IB4, red) for endothelial cells, or neither, following cardiomyocyte isolation, enrichment and purification steps, including the CD31 Ab-bead bound fraction. Scale bar is 100μm. (B) PFA-fixed cardiac cell types were quantified after each step of isolation and purification from C57BL/6J P2 (n=3, ~2000 cells/n), P10 (n=3-4, ~1500 cells/n), P13 (n=3, ~1500 cells/n), or adult (n=3, ~1500 cells/n) hearts using immunocytochemistry (see representative images in Supplementary Fig. S2-5). Cardiomyocytes, CMs (TO-PRO-3^+^, cTnT^+^); endothelial cells, ECs, (TO-PRO-3^+^, IB4^+^); other cells (TO-PRO-3^+^, cTnT^−^, IB4^−^). One-way ANOVA with Dunnett’s multiple comparisons was used to compare cells at each step relative to those after the isolation step. *P<0.05, **P<0.01. ***P<0.001, ****P<0.0001. (C) Representative flow cytometric data evaluating the percentage of nucleated cells (propidium iodide, PI^+^) that were cardiomyocytes (cTnT^+^) or non-myocytes (cTnT^−^) from P2, P10 and P70 PFA-fixed cardiac cell populations after the isolation and purification steps. Approximately 100,000-500,000 cells were analysed per sample and percentages were calculated as a total of the nucleated cell population (PI^+^). The gating strategy was determined per sample based on the controls for the same fractions of the sample that were unstained and single stained (PI^+^ only, cTnT^+^ only). Analysed by unpaired Welch’s t-test. ***P<0.001, ****P<0.0001.

### 3.7 Sex differences identified in the neonatal cardiomyocyte transcriptome

There are surprisingly few studies that have identified sex-specific genes in cardiomyocytes. To evaluate sex differences in neonatal cardiomyocytes, we performed RNA-Seq studies using cardiomyocytes purified from one P2 male and one P2 female heart per litter (n=4 litters, litter sizes of 6-8 pups, Fig. 7A). Of note, traditional “batch chopping” methods for cardiomyocyte isolation would require eight hearts to be pooled for a single replicate. DNA was simultaneously extracted with RNA from the same cardiomyocyte preparation to verify the sex of each mouse pup by PCR amplification of the sex-determining region (SRY on Chr Y, Supplementary Fig. S6). Body weights were similar between litters and sexes (Fig. 7B). Transcriptomic analysis revealed a total of 12,854 genes (> 1cpm in 4 samples) expressed in neonatal cardiomyocytes. Unsupervised clustering of these genes grouped neonatal cardiomyocytes by litter (Fig. 7C), indicating greater variation between biological litters. Differential expression analysis between sexes revealed nine differentially expressed genes (DEGs, FDR<0.05; Fig. 7D, E), all located on the sex chromosomes (Chr Y: *Gm29650*, *Eif2s3y*, *Ddx3y*, *Uty*, *Kdm5d;* Chr X: *Kdm5c*, *Kdm6a*, *Xist*, *Eif2s3x*).

**Fig. 7.**
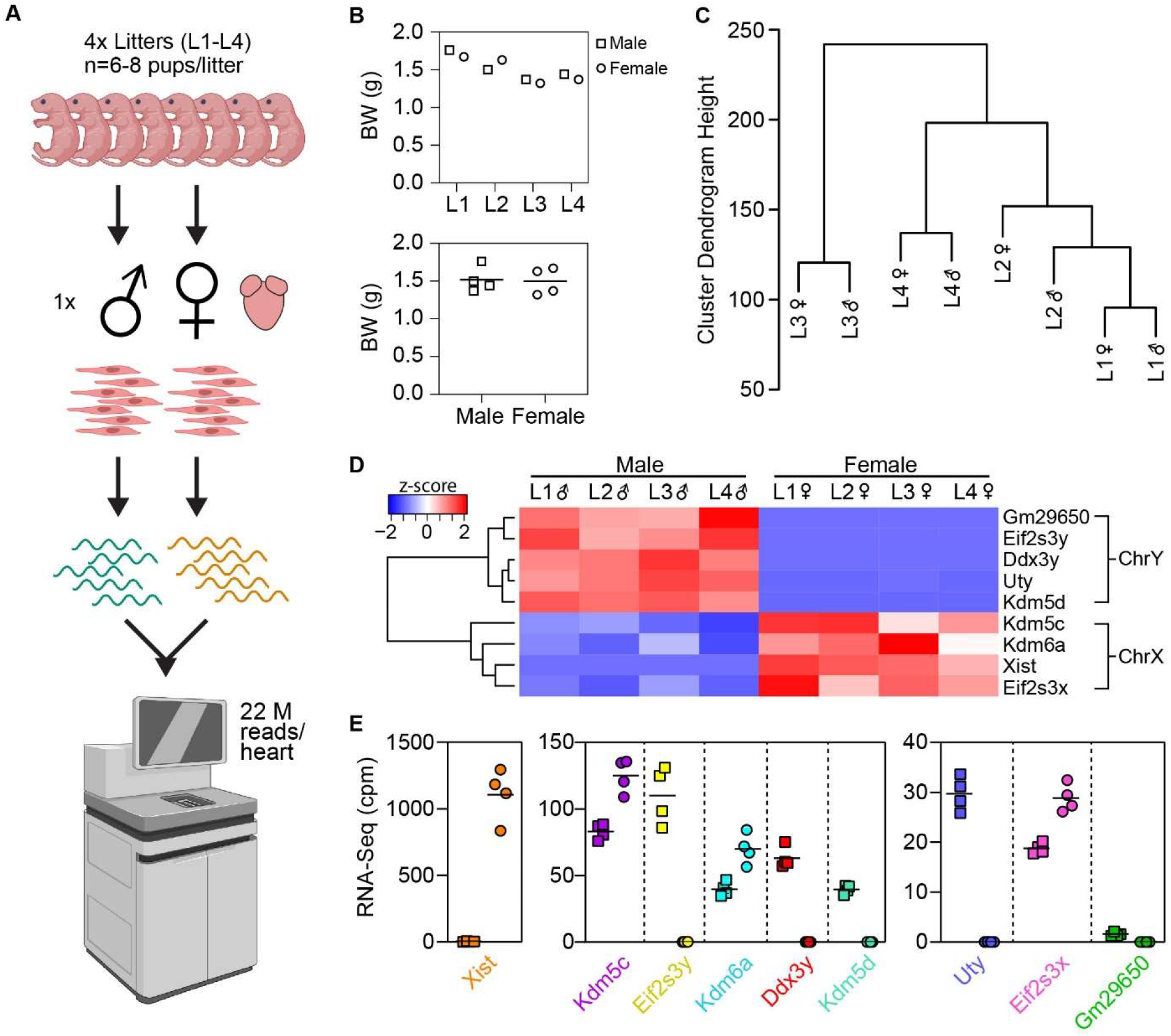
RNA-Seq analysis of neonatal cardiomyocytes reveals hierarchical clustering of litters and sex-specific genes. (A) Schematic of the experimental design was created with BioRender.com. Cardiomyocytes were purified from one P2 male and female heart per litter of 6-8 C57BL/6J mouse pups (x4 litters, L1-L4). Poly(A) RNA extracted from cardiomyocytes and sequenced using NovaSeq 6000 sequencing platform generating ~22M reads per heart. (B) Body weights (BW) of male (□) and female (○) littermates (n=4 per sex). (C) Hierarchical clustering dendrogram (Ward method) based on the expression of 12,854 genes (>1 count per million, in four samples). (D) Heatmap of nine differentially expressed genes (DEGs) between male and female cardiomyocyte samples (n=4 per sex) were identified by edge R (FDR<0.05) and encoded on chromosome Y (ChrY) and chromosome X (ChrX). (E) Abundance of the nine DEGs (cpm, counts per million) presented as dot plots with mean, ordered highest (left) to lowest (right) counts in male (□) and female (○) cardiomyocytes.

## 4 Discussion

We have developed methods for the morphological and molecular study of high-quality cardiomyocytes from individual murine hearts of any postnatal age. This has been achieved by establishing (1) a standardised cardiomyocyte isolation method using Langendorff perfusion, (2) a simple immunomagnetic cell separation step to purify cardiomyocytes, and (3) a cardiomyocyte *in situ* fixation technique that preserves cytoarchitecture. In doing so, cardiomyocyte morphology was profiled at different stages of postnatal development with respect to cell size and nucleation status, including proliferating neonatal cardiomyocytes. We also uncovered sex differences in the neonatal cardiomyocyte transcriptome that have not been previously described.

For successful cardiomyocyte isolations from neonatal and infant hearts, our *in situ* cannulation method was essential for Langendorff retrograde perfusion building upon our previous work[21]. To the best of our knowledge, only one other group has described the isolation of neonatal cardiomyocytes by Langendorff retrograde perfusion[22], and this involved complex *ex vivo* microscopic manipulations to fix a glass cannula to the ascending aorta with an epoxy. More recently, to overcome the technical difficulties associated with cannulating the aorta of younger mice, the same investigators performed antegrade perfusion via a needle inserted into the apex of the left ventricle and clamped the aorta[23]; an approach developed for the isolation of adult left ventricular cardiomyocytes[24]. Although Langendorff perfusions by *ex vivo* aortic cannulations are challenging[15,16,25], our *in situ* aortic cannulation technique is easier to perform, not only in adult hearts, but especially in younger hearts. This could be useful for other Langendorff-based experiments, such as establishing working heart models in neonates and infants, which was recently described for P10 hearts[26].

Another key advance was the optimisation of tissue digestion conditions to isolate cardiomyocytes in a comparable time (~13 min) across hearts of all ages. This standardisation of cardiomyocyte isolations is useful for age-related experiments. Yield is one of the most important measures for successful cardiomyocyte isolations, and yet, it is not widely reported. One commercial neonatal cardiomyocyte isolation kit (Pierce primary cardiomyocyte isolation kit, Thermo Scientific) is reported to yield ~2 million cardiomyocytes per 100 mg of tissue using ~10-15 murine hearts per experiment. In contrast, our cardiomyocyte yield is ~11-fold higher, that is, ~1.56 million ventricular cardiomyocytes per neonatal heart (~7 mg), which is equivalent to ~22 million cardiomyocytes per 100 mg of tissue. Others using the traditional “batch chopping” method reported an extrapolated yield of ~0.6×10^6^ per heart[14,27], which is also ~2.6-fold lower than those achieved using our method. Even improved “batch chopping” methods that have a similar yield to ours (~1.3×10^6^ extrapolated per heart) still required multiple neonatal hearts and take >12-16 h per isolation[27]. Moreover, we found neonatal and infant cardiomyocytes (rod-shaped) had better structural integrity than those isolated by antegrade perfusion since the proportion of rodshaped cells was higher by 20-30%[23]. In terms of adult ventricular cardiomyocytes our yield of 2.2 million cells per heart with a viability of ~72% is comparable to that of other Langendorff perfusion methods[23,28,29]. Together, the combination of *in situ* aortic cannulation and Langendorff digestion enables the isolation of high-quality cardiomyocytes in high yield from individual hearts of any postnatal age, i.e., without using age-dependent procedures or pooling hearts. This is advantageous as it provides individual replicates and reduces animal usage.

We have also shown that cardiomyocytes fixed *in situ* immediately after tissue digestion retained cell integrity; their cytoarchitecture closely resembling that seen *in vivo* with most cells (~94%) remaining intact rod- or spindle-shaped. This suggests that the process of Langendorff tissue digestion itself is gentle and not responsible for the loss of cell structure or viability, which occurs during subsequent cell dissociation steps. When evaluating cardiomyocyte size and nucleation state, as well as cardiac cell populations, it is preferable to isolate cardiomyocytes to obviate the inherent difficulties of using tissue sections[30–33]. These difficulties include the inability to visualise whole cells in a section (even in thick sections) because cardiomyocytes, particularly those in adult hearts, are large and frequently overlapping, irregularly shaped, and arranged in various orientations. Hence, we used *in situ* fixed cardiomyocytes to evaluate cell area by planimetry, which given the irregularities of cell outlines provides more accurate determinations of cell area than calculations based on cell length and width. We have used planimetry previously to demonstrate cellular hypertrophy after cardiac injury[18,19]. Planimetry determinations of cell area revealed that during postnatal development cardiomyocytes increase in size commensurately with cardiac growth, indicating that cellular growth is the main driver of heart growth. In addition, the binucleation index of infant cardiomyocytes at P13 (81%) was close to that in adult cells (85%), albeit lower than previous reported assessments of 98% and 95% at P8[34] and P10[35], respectively. This may reflect differences in cell isolation methods, mouse strain, or litter size, or housing conditions, which, in turn, influence the rate of cardiac growth. Importantly, *in situ* fixation captured spindle-shaped neonatal cells undergoing cytokinesis; an observation only demonstrated previously in tissue sections[36] or rounded cells. Thus, *in situ* fixation is a valuable method for accurately studying cardiomyocyte morphology as well as for the detection and spatial localisation of proteins of interest, such as Aurora B kinase, to better appreciate cell function.

Immunocytochemistry analysis indicated cardiomyocytes were the predominant cell type in neonatal (58%) but not infant cardiac ventricles (36%); cardiomyocytes averaged 51% in adult males, which is similar to that reported for adult humans[38]. It is of interest that in agreement with recent adult human[38] and mouse studies[32], but in contrast to previous reports[35,37], we found endothelial cells were the main non-myocyte cell type in adult hearts; a finding we also observed in neonatal and infant hearts (Fig 6B). A caveat for this observation is the incomplete digestion of cardiac ventricles, as evidenced from endothelial cells still being found in undigested clumps or as strings of cells. This is unsurprising given that harsher digestion conditions (e.g., incubation of chopped tissue with collagenase A[39] or collagenase IV, similar to collagenase D, and dispase II[32] for 45-60 min), are usually required to isolate endothelial cells. It may be possible to develop future protocols that combine Langendorf perfusion, which is gentle, with the addition of other extracellular matrix-targeting enzymes for effective isolation of all cardiac cell types from an individual heart.

Although non-myocytes were partially removed by differential centrifugation [16], significant endothelial cell contamination remained in infant and adult preparations. This varied between tissue digestions and batches of enzymes. For this reason, contaminating endothelial cells were selectively removed by immunomagnetic cell separation whilst neonatal cardiomyocytes were purified by a similar method using a kit targeting the removal of various non-myocytes. Importantly, these purification steps were rapidly completed across all ages (22 min), i.e., without hours of cell culture, and consistently resulted in highly purified (~95%) and viable cardiomyocytes. In these studies, purity after the final step was evaluated by immunocytochemistry and flow cytometry, and further validated by qRT-PCR. For our purposes, we found immunocytochemistry to be the most reliable method for quantifying cell populations. One of the technical limitations with flow cytometric evaluations highlighted by our study was the requirement of detergent for intracellular antibody staining, which hinders efficient cell pelleting in the following wash steps and leads to cell loss. This is likely to bias the analyses towards cells with a higher density. Despite this, up to half a million cells were analysed under the same experimental conditions and there was a significant reduction in the non-myocyte population post-purification.

The estimated yields of cardiomyocyte DNA, RNA, and protein obtained from individual hearts using our isolation and purification method are sufficient for cardiomyocyte biology studies involving omics-based experiments (genomic, epigenomic, transcriptomic, and proteomic studies) or biochemical assays, such as PCR, qRT-PCR, immunocytochemistry and Western blot. As shown here in our RNA-Seq studies of purified cardiomyocytes from individual male and female neonatal hearts, we determined sex-specific genes in neonatal cardiomyocytes. Our data revealed that gene expression profiles clustered predominantly based on litter origin rather than sex. Despite this, nine sex-specific genes were identified that were not only different between males and females but also restricted to X and Y chromosomes. One gene well-known to be sex-specific across many cell types, *Xist*, a long non-coding RNA required for X inactivation, showed the highest expression (>1000 fold-change) in females. This validates *Xist* as a useful marker for sex identification of female cardiomyocytes. Three of the nine genes that we found to be sex-linked (*Uty/Kdm6c*, *Xist*, *Kdm5d/Jarid1d*) have been identified previously in a study of single (cardiomyocyte) nuclei from adult human ventricles[40]. In addition to these genes, an earlier study also identified *Eif2s3y* and *Dddx3y* in the adult mouse and human myocardium[41]. This suggests that expression of these sex-specific genes is maintained from birth to adulthood and that the expression of *Gm29650, Kdm5c/Jarid1c* (a paralog of *Kdm5d*), *Kdm6a/Utx* (a paralog of *Uty*), and *Eif2s3x* may be unique to the neonatal period. Overall, cardiomyocytes purified by our method can be used for a wide range of assays to address cell-type specific questions, and cardiomyocytes from younger hearts, like adult hearts, can be explored on an individual replicate basis to evaluate individual genetic, therapeutic, surgical, age, sex, and biological differences.

In summary, we have established a rapid procedure to isolate and purify cardiomyocytes in high yield, viability, and purity from murine hearts at any postnatal age. This method is useful for age-related studies in cardiomyocytes and for the analysis of individual biological replicates, particularly in neonatal and infant mice. Indeed, our data are the first to identify sex differences in the neonatal cardiomyocyte transcriptome. Moreover, Langendorff perfusion combined with *in situ* fixation allows preservation of the native cytoarchitecture for accurately assessing cardiomyocyte morphology and proliferative status. These procedures represent a significant advance in the accessibility and application of primary cardiomyocytes as a model system important for studying cell biology, postnatal heart development, cardiomyocyte regenerative mechanisms and treatments for cardiovascular diseases.

## Author contributions

AMN, SRH, AYC, ML and MT designed research, conducted experiments, analysed data; PEY, DTH, AMN analysed RNA-Seq data; NN and AH conceived research; NJS designed research; AMN, SEI and RMG conceived research, interpreted the data and wrote the manuscript; all authors reviewed the manuscript.

## Acknowledgements

We gratefully acknowledge the Micro Imaging Facility, The Victor Chang Cardiac Research Institute Innovation Centre for immunofluorescence imaging and The Garvan-Weizmann Centre for Cellular Genomics for assistance with flow cytometry studies. We also acknowledge the services and facilities of Australian Genome Research Facility (AGRF) for generating RNA-Sequencing data. Schematics shown in the graphical abstract and Fig. 7A were created with BioRender.com.

