## Supplementary material for "Standardised method for cardiomyocyte isolation and purification from individual murine neonatal, infant, and adult hearts"

Short title: Isolating cardiomyocytes in postnatal development

---

[1] Present address: [1]Cardiac Regeneration Research Institute, Wenzhou Medical University, Wenzhou, 325035, China

### SUPPLEMENTARY METHODS

#### Langendorff retrograde perfusion

Prior to cannulation, the heat exchanger and perfusate line were rinsed with perfusion buffer. The volume capacity of the rig is 10 mL, which is primed with perfusion buffer (~4-9 mL depending on age, Table 1) and the remaining volume filled with digestion buffer ensuring no air bubbles were trapped inside the apparatus. Buffers were kept at RT outside the water bath. Immediately before mounting the heart, valve 2 (at the outflow nozzle; Figure 1B) was opened and the pump was turned on to commence perfusion. The pre-heated water-jacketed organ bath was moved upwards to surround the heart and maintain it in a warm environment. To avoid air bubbles entering the heart, perfusion was stopped prior to the last of the digestion buffer entering the heart. The total time taken to perfuse hearts at any postnatal age was <14 min (Table 1). Atria were removed while the heart was still mounted and then the heart was transected at the atrioventricular junction to release the ventricles, which were collected into a petri dish (60 x 15 mm, cat.#:351007, Corning) containing transfer buffer (~1- 2 mL; Supplementary material online, Table S2). Note: with two Langendorff rigs, one person can sequentially cannulate, mount, and perfuse two hearts, enabling cardiomyocyte isolations and purifications from eight hearts in one day.

#### Tissue digestion of hearts from neonates through to adults

The concentration of proteolytic enzymes, volume of buffers and perfusate flow rate for tissue digestion on the Langendorff apparatus were determined empirically (see Table 1 and Supplementary material online, Tables S2, S3). Three separate enzyme stock solutions, based on Liao and Jain (2007)[1], sufficient for the total number of hearts to be digested on a given day, were prepared freshly in perfusion buffer (1 ml). The stock concentrations of proteolytic enzymes: collagenase B (0.3 mg/g body weight), collagenase D (0.4 mg/g body weight) and protease XIV (0.05mg/g body weight), were calculated using the body weight of adult mice. However, for neonates and infants stock solutions based on actual body weights did not provide high enough enzyme concentrations to allow efficient digestion, so empirical adjustments to weights of 5 and 8 g, respectively, were required. Stock solutions were diluted in 16-30 mL of perfusion buffer, referred to as “digestion buffer” (Table 1), and this was used to digest hearts. An example of the calculation for preparing digestion buffer for six neonatal hearts is shown in Supplementary material online, Table S3. Yield, cell viability and structural integrity (i.e., rod-shaped cardiomyocytes) are dependent on the efficiency of tissue digestion, with under-digestion leading to clumps of tissue that are either removed by filtration thereby reducing yield, or that require more vigorous tissue disaggregation resulting in shear stress and cell death.

#### Cardiomyocyte isolation

Digested ventricles were teased apart with fine scissors and forceps into ~10-15 pieces and dissociated into a cell suspension by gentle trituration (15 times) using a transfer pipette. The cut tip of a P1000 pipette (for neonatal hearts) or transfer pipettes (for infant and adult hearts; cat. #: 222 and 202, Samco Scientific) trimmed to create a wide bore to minimise cell shearing. Tissue and cells were transferred from the petri dish into a 15 mL Falcon tube, and the dish was washed with transfer buffer to collect any remaining material, which was added to the tube. After further gentle trituration with the gradual addition of transfer buffer (up to 6, 8, or 10 mL for neonatal, infant or adult cell preparations, respectively), the cell suspension was passed through a 70 µm mesh filter (cat. #: 352350, Corning) for neonatal and 200 μm filter (cat. #: 340644, Corning) for infant and adult cardiomyocytes to remove any undigested tissue aggregates (summarised in Table 1). Cells were uniformly resuspended with a wide bore transfer pipette by gentle trituration and immediately loaded onto a haemocytometer (Improved Neubauer counting chamber, AC100, Hawksley) for determination of cardiomyocyte number and the proportion of viable rod-shaped cardiomyocytes. Only infant and adult cell preparations with >60% of cells being rod-shaped, were used. Viability was determined either by Trypan blue exclusion (neonatal cardiomyocyte) or by morphology (infant and adult cardiomyocytes): viable cells being rod-shaped and excluding Trypan blue, whereas round cells were non-viable and took up Trypan blue. Note: Cardiomyocyte cell counts using a haemocytometer were interpreted as estimates. Cardiomyocytes that intersected the perimeter of the haemocytometer square but mostly lay inside were included in the cell count because they are long and irregularly shaped. Neonatal cardiomyocytes can be readily distinguished as spindle-shaped cells, whereas rounded neonatal cells can easily be distinguished by the granulated appearance of their myofibrils.

### Cardiomyocyte enrichment

Infant and adult cardiomyocytes were enriched by low-speed centrifugation (3 x 3 min, 55 *g* or 20 *g*, respectively), without addition of $Ca^{2+}$, to separate them from the smaller non-myocytes. The resulting cardiomyocyte-enriched pellet was resuspended in 6 or 8 mL of transfer buffer for infant or adult cells, respectively, by gentle trituration using a transfer pipette. After enrichment, cardiomyocyte number and the proportion of viable rod-shaped cardiomyocytes were estimated by haemocytometer counting.

### Tissue collection and histology

Litters of 6-8 mice were sacrificed using age-appropriate methods (Table 1) and hearts were harvested, rinsed in saline, and blotted dry before being weighed. Adult male mice were sacrificed by $CO_2$ inhalation. For histology purposes, freshly dissected aortae or ventricles were rinsed with saline to remove blood, fixed (2% PFA, 4 hr, RT), washed in PBS and cryoprotected in sucrose. Tissues were embedded in OCT and sectioned (7 μm, Leica CM 1950). Sections were stained with H&E. Images for Figure 1C were taken with Leica DM6000B light microscope (power mosaic at 20x objective).

### Immunocytochemistry

Fixed cells were mounted onto slides by Cytospin centrifugation (500 rpm, 5 min), washed in PBS (3x 5 min), incubated with blocking buffer (3% BSA in PBS; 1 hr, RT) followed by mouse anti-cardiac troponin T antibody (cTnT, cat. #: ab10214, Abcam, 1:600 in blocking buffer; overnight, 4°C), and then washed in PBS (x3 5 min). The following steps were performed in the dark. Cells were incubated with goat anti-mouse IgG Alexa Fluor 488 (1:1000) in PBS with 1% BSA (1 hr, RT), washed twice in PBS, twice in lectin buffer (50 mM Tris base, 0.87% NaCl, 0.02% $MgCl_2$, 0.01% $CaCl_2$ pH 7.6 with 0.01% Tween-20; 5 min each, RT). Slides were incubated with an endothelial cell marker (Isolectin GS-IB4-568 conjugate; cat. #: I21412, Life Technologies, 1:100 in lectin buffer with 3% BSA, 1 hr, RT), followed by three washes in lectin buffer before DNA staining (TO-PRO-3; cat. #: T3605, Life Technologies, 1:1000 of 1 mg/mL) in lectin buffer (15 min, RT). Slides were washed three times in lectin buffer (5 min) before mounting with PVA-DABCO mounting medium and imaged on a Zeiss Axio Observer Z1 (inverted) equipped with a LSM 700 confocal scan head (tile scan: 10 or 15 fields, z-stack of >6 planes, 25x or 40x magnification). Laminin staining was performed as previously described[2].

For Aurora B kinase staining, slides were washed twice in PBS before performing HIER (Vector, H-3300, 95°C, 5 min) and the solution was allowed to cool for 1 hr. Following two washes of PBS, endogenous peroxidase activity was quenched (0.3% $H_2O_2$ in 70% methanol in PBS; 30min, RT). Slides were washed three times and incubated with blocking buffer (3% normal donkey serum, 3% BSA and 0.1% triton X-100 in PBS; 1 hr, RT) followed by mouse anti-cardiac troponin T antibody (cTnT, cat. #: ab10214, Abcam, 1:600) and rabbit anti-aurora B antibody (cat. #: ab2254, Abcam, 1:50) in blocking buffer overnight at 4°C, then washed four times in PBS. The following steps were performed in the dark. Cells were incubated with donkey anti-mouse IgG Alexa Fluor 488 (1:1000) and biotinylated donkey anti-rabbit IgG (Jackson ImmunoResearch, cat. #: 711-065-152, 1:250) in PBS with 1% BSA; 1 hr, RT), washed four times in PBS, incubated with avidin biotin complex (Vector Laboratories, Vectorstain Elite ABC-HRP kit, cat. #: PK-6101, reagent A 1:100, reagent B 1:100 in PBS; 1 hr, RT), washed twice in PBS, incubated with TSA-cy3 (Perkin Elmer, cat. #: NEL704A001KT, 1:100 in amplification diluent; 3 min, RT) followed by three washes in PBS before DNA staining (TO-PRO-3; cat. #: T3605, Life Technologies, 1:1000 of 1mg/mL, 15 min, RT). Slides were washed three times in PBS before mounting with PVA-DABCO mounting medium and imaged on a Zeiss Axio Imager Z1 (upright) equipped with a LSM 700 confocal scan head.

**Quantifying cardiac cell populations, cardiomyocyte area and nucleation status**

Cell populations and nucleation state were quantified using the multi-point tool in ImageJ software (v1.53e). All nuclei were counted, and cells were categorised as cardiomyocytes (cTnT+), endothelial cells (IB4+) or "other" (cTnT- and IB4-). Cardiomyocyte area and nuclei state (mononucleated, binucleated or multinucleated) were measured using *in situ* fixed cardiomyocytes. Overlapping cardiomyocytes were excluded from analysis. Cardiomyocyte planimetry area was determined using ImageJ with automated outlining of cells and area calculated relative to the scale bar. The number of nuclei per cell was counted using ImageJ. Cells with two overlapping nuclei, possibly undergoing karyokinesis, were counted as mononucleated cardiomyocytes.

**Flow cytometry**

Cardiac cells were fixed either pre-enrichment or post-purification, washed (2000 *g* for 3 min) and incubated with BD Perm/Wash buffer for 10 min at RT. Cells were stained in the dark with Alexa Fluor 647 anti-cardiac troponin T, cTnT (#: 565744; 1:200 dilution, 30 min, 4$^{o}$C) followed by two washes, and then resuspended in PBS and 0.5% BSA in FACS tubes (100 µl). Propidium iodide (PI; 1:100; 1 mg/mL) was added 10 min prior to running samples on a BD FACS Canto II Cell Analyzer cytometer fitted with a 100 µm nozzle. Larger adult cardiomyocytes were resuspended in a greater volume (>300 µl) and maintained in suspension by frequently flicking the tube to prevent them settling during flow cytometry. Flow samples were run at a low or medium flow rate until the tube was completely emptied to prevent cell bias due to larger cells settling more rapidly compared to smaller cells. Cardiac cells from one heart were separated into three controls tubes (unstained, cTnT only stained, PI only stained) and one sample tube (cTnT and PI stained) to determine the gating strategy for analyses using FlowJo software. Note: Cell permeabilisation was required for intracellular staining for flow cytometry. During this step, cell density decreases resulting in a looser cell pellet that leads to cell losses in the following wash steps[3], particularly smaller cells. Cell loss occurred despite testing different detergents and concentrations as well as increasing the centrifugal force for washes. Cell permeabilization for immunocytochemistry was performed when cells were secured onto slides resulting in little or no cell loss. Thus, flow cytometric analyses may lead to an overestimation of the cardiomyocyte population.

**RNA extraction and qRT-PCR**

Fractions of cells were collected at various steps throughout the P2, P10, and adult cardiomyocyte isolation and purification procedure. Samples were homogenised (with a 5x75 mm generator for P2 and P10, and a 7x95 mm generator for P70, PRO Scientific homogenizer) in QIAzol Lysis Reagent (700 µL) on ice and RNA extracted following the manufacturer's instructions (miRNeasy Mini Kit, cat. #: 217004, Qiagen Sciences). For cDNA synthesis, 500 ng of RNA was used as input for DNase treatment and reverse transcription (SuperScript IV VILO Master Mix with ezDNase Enzyme kit, Invitrogen). qRT-PCR was performed with diluted cDNA (1:30) using a TaqMan probes (TaqMan Gene Expression Master Mix, Thermo Fisher; Table S5).

**DNA extraction and yield**

DNA was extracted using the DNeasy Blood & Tissue Kit (cat. #: 69506, Qiagen) and animal tissues protocol (modified as follows). Adult cardiomyocytes samples were re-suspended in ATL buffer (360 µL), homogenised and sonicated, followed by the addition of proteinase K (40 µL) and incubated for 3 hr at 56°C. Neonatal and infant cardiomyocyte samples were re-suspended in ATL buffer (180 µL) and sonicated, and then proteinase K (20 µl) was added and samples were also incubated for 3 hr at 56°C. Cardiomyocyte DNA yields were measured by a Qubit 2.0 fluorometer and Qubit dsDNA High Sensitivity assay.

**Protein extraction and yield**

Cardiomyocyte samples were re-suspended in ice-cold lysis buffer (1x RIPA buffer, cat #: 9806 and 1 mM PMSF) with protease (cat #: 11836170001, Roche) and phosphatase inhibitors (cat #: P0044 and P5726, Sigma-Aldrich), homogenised and sonicated, and incubated for 45 min at 4°C on gentle agitation. Samples were centrifuged at 16,000 *g* for 15 min at 4°C and the protein lysate supernatant collected. Protein yields were measured using a BCA assay (Pierce BCA Protein Assay Kit, cat #: 23227).

**DNA extraction for SRY PCR**

RNA and DNA were extracted from the same samples used for RNA-Sequencing. RNA was removed in the upper aqueous layer upon phase separation after the addition of chloroform and centrifugation. DNA lies at the interphase in between the upper RNA layer and lower organic phenol phase containing protein. DNA was extracted during phase separation using the user-developed protocol provided by Qiagen resources online. Briefly, the interphase and lower phase was mixed with 100% ethanol and centrifuged to sediment DNA, which was washed with sodium citrate three times and then 75% ethanol before air-drying the DNA pellet. DNA was redissolved in sodium hydroxide (8mM) and Tris-EDTA (1 mM), and stored at 4°C.

To confirm the sex of samples collected for RNA-Sequencing, DNA (8-10ng) was PCR amplified using primers[4] for the sex determining region Y (SRY) gene and DreamTaq Green Master Mix (cat. #: K1081, Thermo Scientific) with the following PCR conditions: 95°C for 1 min, 26 x cycles of 95°C for 30 sec, 60°C for 30 sec, 72°C for 30 sec, and a final step of 72°C for 10 min. PCR products and 100bp DNA ladder (G210A, Promega) were run on a 2.5% agarose gel in TAE buffer (40 mM Tris Acetate, 1.15% Glacial Acetic Acid, 10 mM EDTA) with ethidium bromide (1 µg/mL of gel) at 100V for 40 min.

**RNA-Sequencing of neonatal cardiomyocytes**

Sequencing libraries were constructed from ~1 µg of total RNA using stranded TruSeq total RNA library preparation kit (Illumina) through a sequencing service provider (Australian Genome Research Facility, Melbourne, Australia). Libraries were sequenced on an Illumina NovaSeq 6000 aiming for a depth of 20 million 1x100bp single end reads per sample. Sequencing reads were trimmed and adaptors using Trimmomatic[5] using parameters LEADING:3 TRAILING:3 SLIDINGWINDOW:4:15 MINLEN:33. Sequence reads were mapped to the mm10 mouse reference genome using RNA STAR (v2.7.3) following the two-pass alignment protocol[6]. Gene expression was quantified from read counts calculated by RNA STAR. Cluster dendrogram (hclust, Ward method) and heatmap of DEGs were all generated via scripts in R. Differential expression analysis was performed using edgeR[7] using log-transformed data (+0.01 to genes with 0 cpm).

### SUPPLEMENTARY FIGURES

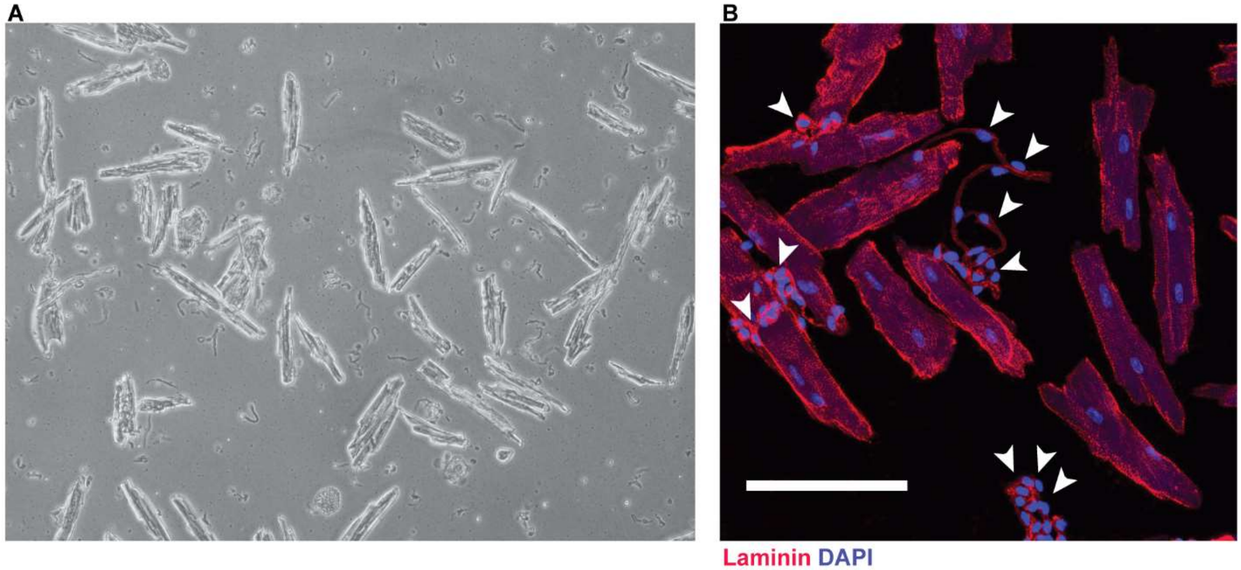

**Supplementary Figure S1: Isolated and enriched adult cardiomyocyte fractions.**

(A) Brightfield image of isolated adult cardiac cells, including non-myocytes, released by enzymatic digestion using the digestion buffer detailed by Liao and Jain (2007)[1]. (B) Immunofluorescence image of adult cardiomyocytes fixed after the enrichment step of differential centrifugation using the digestion buffer detailed by O'Connell *et al.* (2007)[8]. Cell membranes were stained with laminin (red) and DNA with DAPI (blue). Endothelial cells appear as undigested strings and clumps (white arrows) attached to cardiomyocytes. Scale bar is 100 μm.

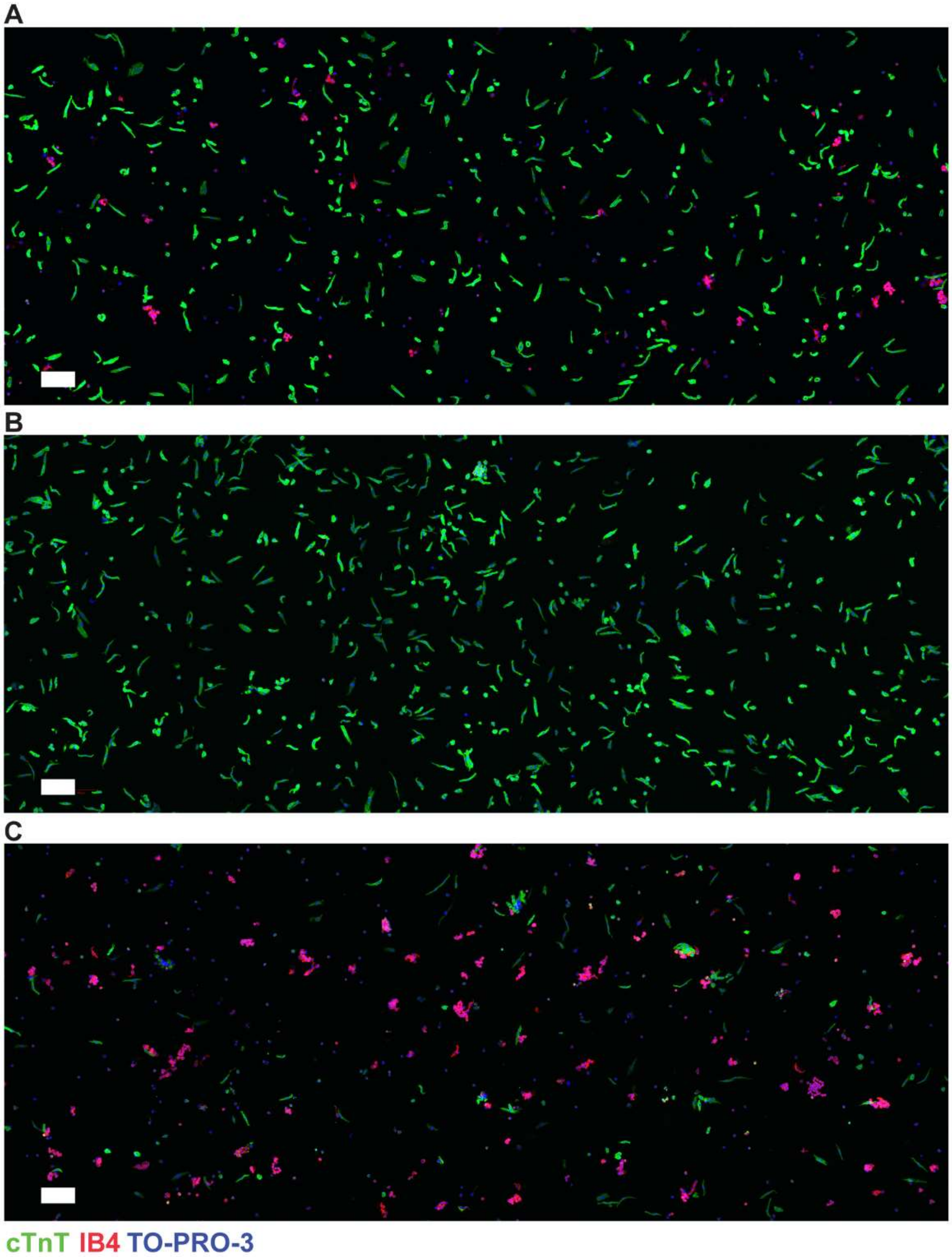

**Supplementary Figure S2: Isolation and purification of neonatal cardiomyocytes.** Immunofluorescence images of cells from a C57BL/6J postnatal day (P) 2 heart fixed after the cardiomyocyte isolation (A) and purification step (B), and in the Ab-bead-bound fraction (C) (10 fields of view per image at 25x), stained for cardiomyocytes (cTnT, green), endothelial cells (IB4, red), and cells with nuclei (TO- PRO-3, blue) but without cTnT or IB4, were quantified (Figure 4B). Scale bar is 100 µm.

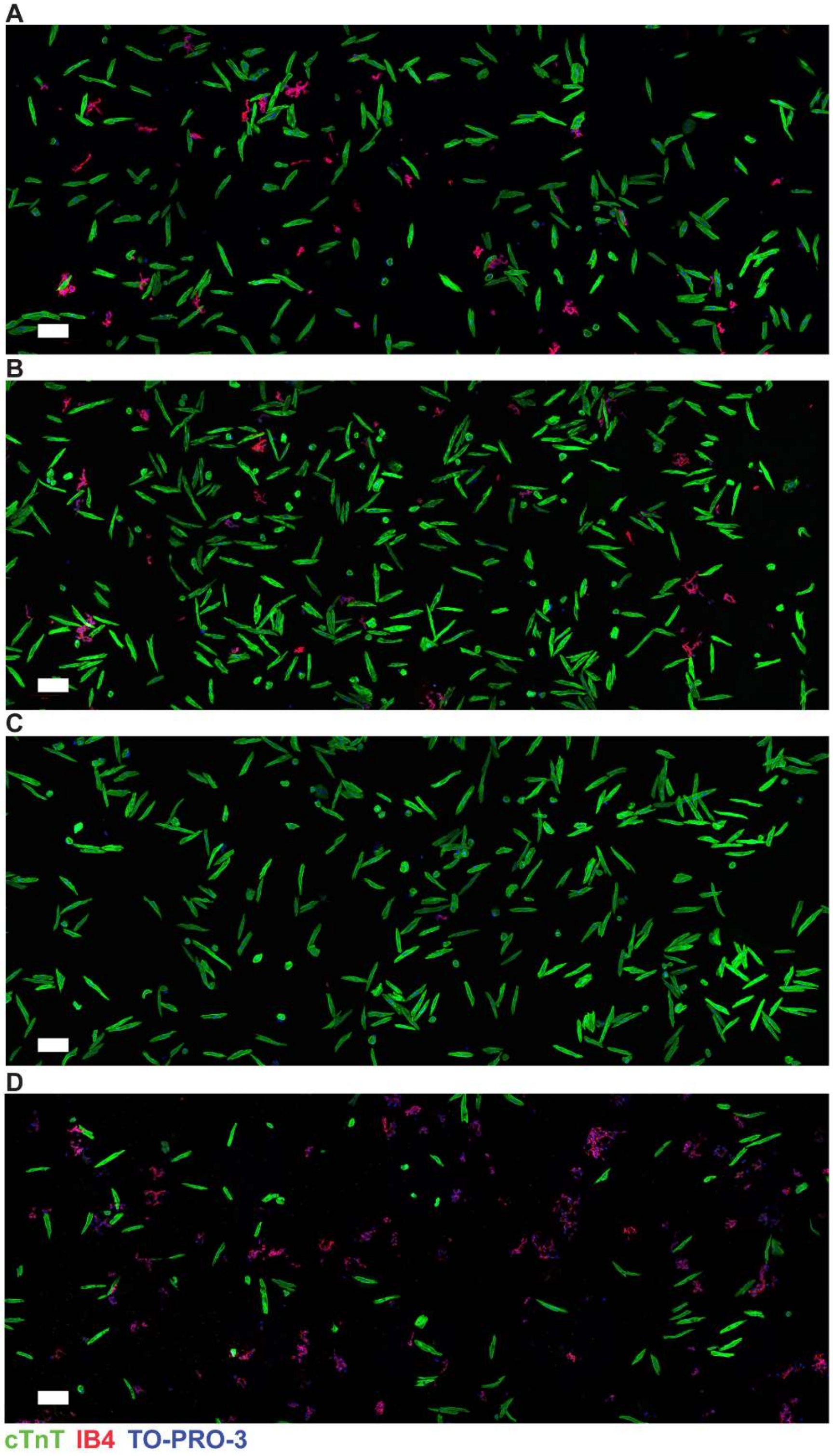

**Supplementary Figure S3: Isolation, enrichment and purification of infant cardiomyocytes.** Immunofluorescence images of cells from a C57BL/6J postnatal day (P) 10 heart fixed after the cardiomyocyte isolation (A), enrichment (B), and purification step (C), and in the Ab-bead-bound fraction (D) (10 fields of view per image at 25x), stained for cardiomyocytes (cTnT, green), endothelial cells (IB4, red), and cells with nuclei (TO- PRO-3, blue) but without cTnT or IB4, were quantified (Figure 4B). Scale bar is 100 µm.

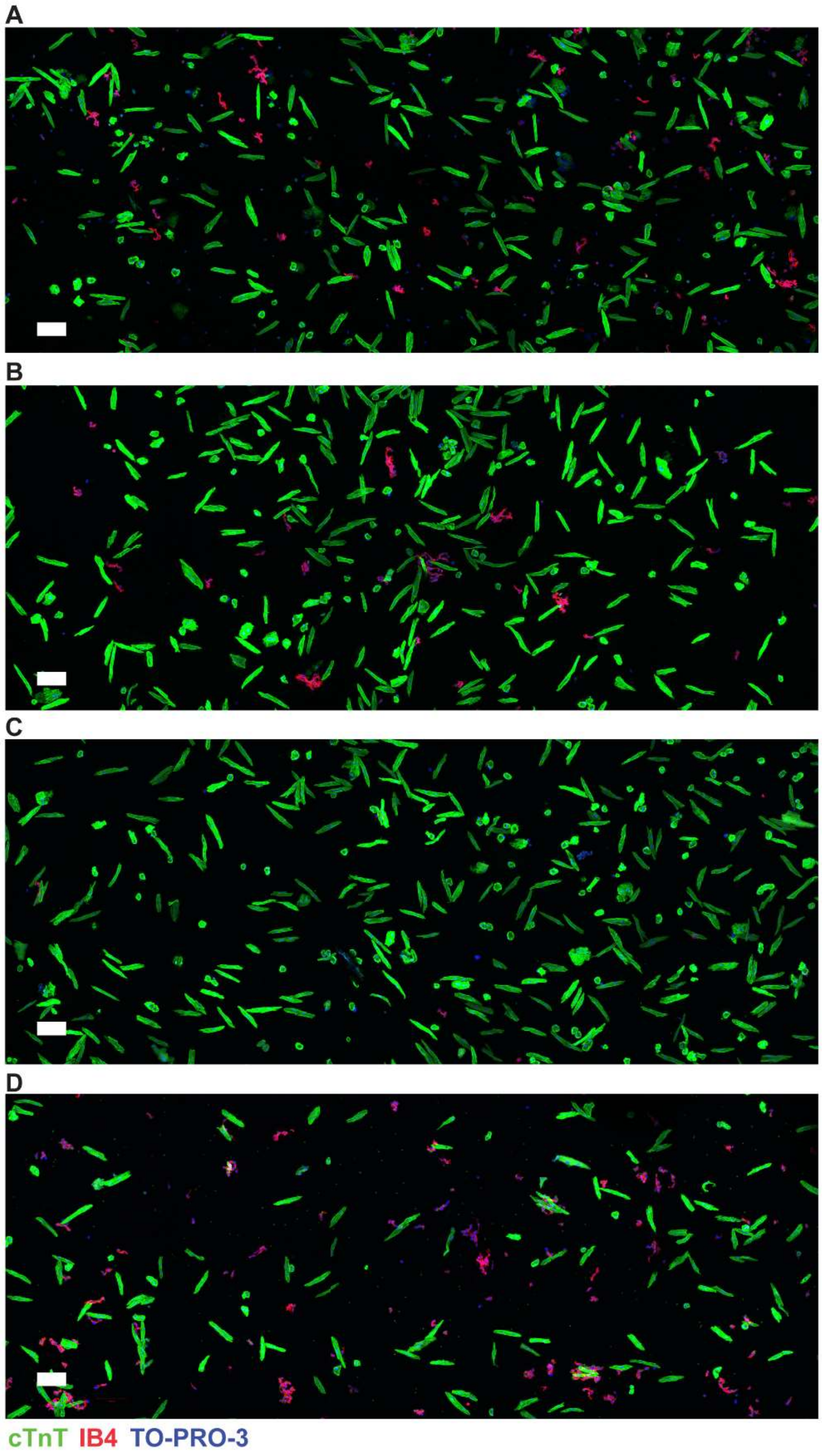

**Supplementary Figure S4: Isolation, enrichment and purification of infant cardiomyocytes.** Immunofluorescence images of cells from a C57BL/6J postnatal day (P) 13 heart fixed after the cardiomyocyte isolation (A), enrichment (B), and purification step (C), and in the Ab-bead-bound fraction (D) (10 fields of view per image at 25x), stained for cardiomyocytes (cTnT, green), endothelial cells (IB4, red), and cells with nuclei (TO- PRO-3, blue) but without cTnT or IB4, were quantified (Figure 4B). Scale bar is 100 µm.

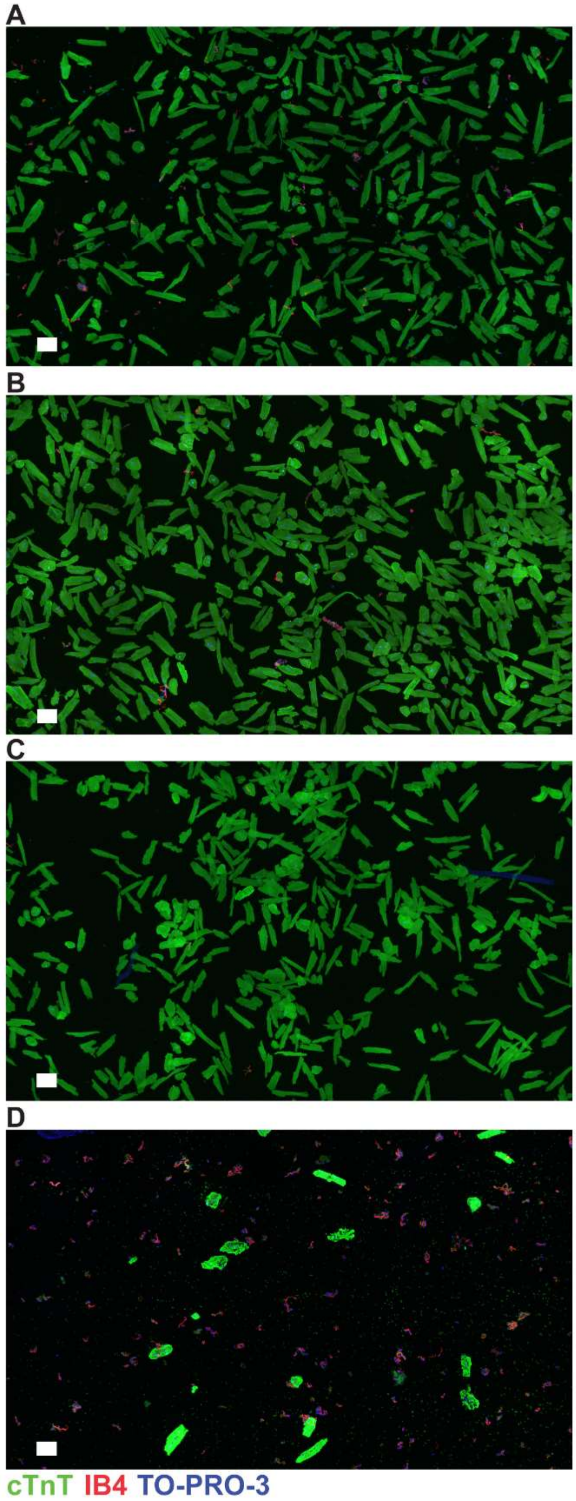

**Supplementary Figure S5: Isolation, enrichment and purification of adult cardiomyocytes.** Immunofluorescence images of cells from a C57BL/6J adult (P) 70 heart fixed after the cardiomyocyte isolation step (A), enrichment step (B) and purification step (C), and in the Ab-bead-bound fraction (D) (15 fields of view per image at 25x), stained for cardiomyocytes (cTnT, green), endothelial cells (IB4, red), and cells with nuclei (TO- PRO-3, blue) but without cTnT or IB4, were quantified (Figure 4B). Scale bar is 100 μm.

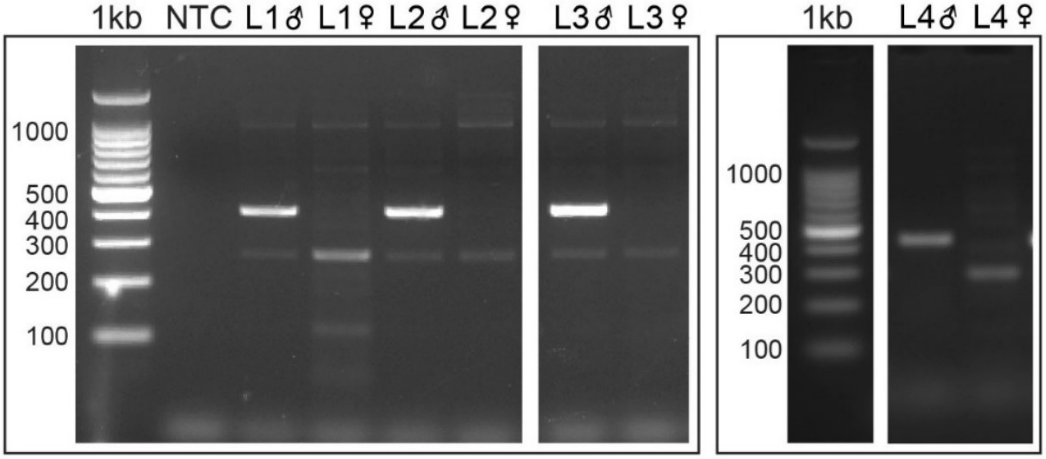

**Supplementary Figure S6: SRY PCR confirmed sex of cardiomyocytes samples.** Agarose gel electrophoresis of sex determining region (SRY) PCR amplicons. Genomic DNA (8-10 ng) from neonatal cardiomyocyte samples were used as template for PCR to confirm the sex of the same samples prior to RNA-Sequencing. The presence of a SRY band at 405bp indicated the sample came from a male. NTC, non-template control has no DNA input in the PCR reaction.

### SUPPLEMENTARY TABLES

**Supplementary Table S1: Materials used for *in situ* cannulation of the heart and cardiomyocyte purification**

| Materials | Company | Cat. # |
|---|---|---|
| **For P2 hearts** | | |
| Fine Scissors ToughCut | Fine Science Tools | 14058-09 |
| Hardened Fine Scissors, curved | Fine Science Tools | 14091-09 |
| Extra Fine Graefe Forceps | Fine Science Tools | 11152-10 |
| 2x Dumont Mini Forceps, Style 5 | Fine Science Tools | 11200-14 |
| Cannula needle (30G) | BD Biosciences | 304000 |
| Syringe 10 mL | BD Biosciences | 302146 |
| Black braided silk 6-0, non-sterile | Surgical Specialities | SP102 |
| MACS MultiStand | Miltenyi Biotec | 36024 |
| OctoMACS Separator | Miltenyi Biotec | 130-042-109 |
| **For P10 hearts** | | |
| Fine Scissors, ToughCut | Fine Science Tools | 14058-09 |
| Student Adson Forceps, Serrated | Fine Science Tools | 91106-12 |
| 2x Student Dumont #7 Forceps | Fine Science Tools | 91197-00 |
| Cannula needle (25G) | Terumo Needle | NN*2519RA |
| Syringe 10 mL | BD Biosciences | 302146 |
| Black braided silk 6-0, non-sterile | Surgical Specialities | SP102 |
| MagnaRack Magnetic Separation Rack | Thermo Fisher | CS15000 |
| **For adult hearts** | | |
| Surgical Scissors Sharp | Fine Science Tools | 14002-13 |
| Fine Scissors, ToughCut | Fine Science Tools | 14058-09 |
| Student Adson Forceps, Serrated | Fine Science Tools | 91106-12 |
| 2x Student Dumont #7 Forceps | Fine Science Tools | 91197-00 |
| Cannula Gavage needle (24G) | Fine Science Tools | 18061-24 |
| Syringe 10 mL | BD Biosciences | 302146 |
| Black braided silk 6-0, non-sterile | Surgical Specialities | SP102 |
| MagnaRack™ Magnetic Separation Rack | Thermo Fisher | CS15000 |
| Multipurpose Thermometer | Physitemp | BAT-10 |

**Supplementary Table S2: Buffer compositions for cardiomyocyte isolation and purification**

| Compound | MW (g/mol) | Stock conc. (M) | Final conc. (mM) | To add (mL or g) | Supplier |
|---|---|---|---|---|---|
| **Perfusion buffer (PB), pH 7.2** | | | | Per litre (L) | |
| NaCl | 58.44 | 5 | 135 | 27 mL | Sigma |
| KCl | 1.98 | 1 | 4 | 4 mL | Sigma |
| $MgCl_2$ | 203.30 | 1 | 1 | 1 mL | Merck |
| $NaHPO_4$ | | 0.6 | 0.33 | 0.55 mL | |
| HEPES | 238.31 | | 10 | 2.38 g | PanReac AppliChem |
| BDM | 101.10 | | 15 | 1.5 g | Sigma |
| Taurine | 125.1 | | 5 | 0.63 g | Sigma |
| Glucose | 180.16 | | 10 | 1.8 g | Sigma |
| **Digestion buffer (DB), pH 7.2** | | | | | |
| Perfusion buffer | | | | | |
| Col B | | | 0.3 mg/g | | Roche |
| Col D | | | 0.4 mg/g | | Roche |
| Protease XIV | | | 0.05 mg/g | | Sigma |
| **Transfer buffer (TB), pH 7.4** | | | | Per 500 mL | |
| NaCl | | 5 | 135 | 13.5 mL | |
| KCl | | 1 | 4 | 2 mL | |
| $MgCl_2$ | | 1 | 1 | 0.5 mL | |
| $NaHPO_4$ | | 0.6 | 0.33 | 0.275 mL | |
| HEPES | | | 10 | 1.19 g | |
| BDM | | | 15 | 0.75 g | |
| BSA | | | 5 g/mL | 2.5 g | |
| Glucose | | | 10 | 0.9 g | |
| **Isolation buffer (IB), pH 7.4** | | | | Per 50 mL | |
| PBS | | | | | |
| EDTA | 372.2 | 0.5 | 2 | 200 µL | Sigma |
| BSA | | | 0.1% | 0.25 g | |

Buffers were made up according to Liao and Jain (2007) protocol[1] (collagenase concentrations were altered slightly from their protocol). All buffers were filter sterilized in a fume hood using a Corning 500 mL Vacuum (cat. #: 431097) after adjusting the pH. MW, molecular weight; BDM, 2,3-butanedione monoxime; Col B & D, collagenase B & D; BSA, bovine serum albumin.

**Supplementary Table S3: Example of digestion buffer preparation to isolate neonatal cardiac cells**

| | Stock conc. calculation (mg/mL) | Working conc. calculation per heart | Vol. of stock enzyme added to perfusion buffer (16 mL) per heart (µL) |
|---|---|---|---|
| No. of mice | 6 | 1 | 1 |
| Set BW (g) | 5 | 5 | 5 |
| Total BW (all animals; g) | 30 | | |
| Col B (0.3mg/g) | = 30 x 0.3= 9 | = 5 x 0.3= 1.5 | = (1.5/9) x 1000 = 167 |
| Col D (0.4mg/g) | = 30 x 0.4= 12 | = 5 x 0.4= 2 | = (2/12) x 1000 = 167 |
| Protease XIV (0.05mg/g) | = 30 x 0.05= 1.5 | = 5 x 0.05= 0.25 | = (0.25/1.5) x 1000 = 167 |

BW, body weight; Col B & D, collagenase B & D.

**Supplementary Table S4: Preparation of Ab (CD31)-beads to purify infant and adult cardiomyocytes**

| Reagents | Stock for 20x adult or 40x infant hearts | Per adult heart | Per infant heart |
|---|---|---|---|
| CD31 purified rat anti-mouse Ab (cat #: 553370, BD Pharmingen, µL) | 200 | 10 | 5 |
| Dynabeads sheep anti-rat IgG (cat #: 11035, Invitrogen, µL) | 500 | 25 | 12.5 |
| Isolation buffer (µL) | 300 | 15 | 7.5 |
| Total volume | 1 mL | 50 µL | 25 µL |

Ab-beads were prepared according to the manufacturer’s instructions using these optimised amounts for infant and adult cardiac cell preparations.

**Supplementary Table S5: TaqMan probes**

| TaqMan gene | Protein | TaqMan probe ID |
|---|---|---|
| *Gapdh* | GAPDH | Mm99999915_g1 |
| *Myh6* | α-MHC | Mm00440359_m1 |
| *Tnnt2* | cTnT | Mm01290256_m1 |
| *Pecam1* | PECAM-1 | Mm01242576_m1 |
| *Vwf* | VWF | Mm00550376_m1 |
| *Ddr2* | DDR2 | Mm00445615_m1 |
| *Col1a1* | COL1A1 | Mm00801666_g1 |

Cardiomyocyte markers: *Myh6*, α-myosin heavy chain, and *Tnnt2*, cardiac Troponin T; Endothelial cell markers: *Pecam1*, Platelet Endothelial Cell Adhesion Molecule 1, and *Vwf*, von Willebrand Factor; Fibroblast markers: *Ddr2*, Discoidin Domain Receptor 2, and *Col1a1*, Collagen Type I Alpha 1 Chain.

**Supplementary Table S6: Cardiac cell marker expression throughout the cardiomyocyte isolation and purification procedure**

| | Isolation | Enrichment | Supernatant | Purification | Ab-beads |
|---|---|---|---|---|---|
| **P2 preparations** | | | | | |
| *Myh6* | 1.00 | NA | NA | 1.12 | 0.41 |
| *Tnnt2* | 1.00 | NA | NA | 1.08 | 0.40 |
| *Pecam1* | 1.00 | NA | NA | 0.004 | 10.07 |
| *Vwf* | 1.00 | NA | NA | 0.001 | 4.29 |
| *Ddr2* | 1.00 | NA | NA | 0.09 | 7.63 |
| *Col1a1* | 1.00 | NA | NA | 0.05 | 8.48 |
| n | 3 | NA | NA | 3 | 3 |
| **P10 preparations** | | | | | |
| *Myh6* | 1.00 | 1.19 | 1.29 | 1.76 | 0.91 |
| *Tnnt* | 1.00 | 1.36 | 1.49 | 1.62 | 1.06 |
| *Pecam1* | 1.00 | 0.43 | 3.22 | 0.06 | 1.29 |
| *Vwf* | 1.00 | 0.49 | 8.84 | 0.03 | 2.38 |
| *Ddr2* | 1.00 | 0.30 | 8.63 | 0.06 | 0.75 |
| *Col1a1* | 1.00 | 0.08 | 10.50 | 0.04 | 0.13 |
| n | 3 | 3 | 3 | 3 | 2 |
| **P70 preparations** | | | | | |
| *Myh6* | 1.00 | 1.18 | 0.74 | 0.99 | 0.93 |
| *Tnnt* | 1.00 | 1.02 | 0.63 | 0.99 | 0.84 |
| *Pecam1* | 1.00 | 0.16 | 11.65 | 0.06 | 1.88 |
| *Vwf* | 1.00 | 0.06 | 13.29 | 0.01 | 1.11 |
| *Ddr2* | 1.00 | 0.05 | 14.88 | 0.03 | 1.24 |
| *Col1a1* | 1.00 | 0.03 | 19.62 | 0.02 | 1.06 |
| n | 5 | 2 | 4 | 5 | 3 |

Genes were normalised to *Gapdh* and fold-change calculated relative to the isolation fraction using the $2^{-\Delta\Delta Ct}$ method. Data presented for isolation, enrichment and purification fractions shown in Figure 5A. Cardiomyocyte markers: *Myh6*, α-myosin heavy chain, and *Tnnt2*, cardiac Troponin T; Endothelial cell markers: *Pecam1*, Platelet Endothelial Cell Adhesion Molecule 1, and *Vwf*, von Willebrand Factor; Fibroblast markers: *Ddr2*, Discoidin Domain Receptor 2, and *Col1a1*, Collagen Type I Alpha 1 Chain.
